# PAMP-triggered Genetic Reprogramming Involves Widespread Alternative Transcription Initiation and an Immediate Transcription Factor Wave

**DOI:** 10.1101/2021.02.16.431463

**Authors:** Axel Thieffry, Jette Bornholdt, Andrea Barghetti, Albin Sandelin, Peter Brodersen

## Abstract

Immune responses triggered by pathogen-associated molecular patterns (PAMPs) are key to pathogen defense, but drivers of the genetic reprogramming required to reach the immune state remain incompletely understood in plants. Here, we report a time-course study of the establishment of PAMP-triggered immunity (PTI) using cap analysis of gene expression (CAGE). Our results show that as much as 15% of all PAMP response genes display alternative transcription initiation. In several cases, use of alternative TSSs may be regulatory as it determines inclusion of target peptides or protein domains, or occurrence of upstream open reading frames (uORFs) in mRNA leader sequences. We also find that 60% of PAMP-response genes respond much earlier than previously thought. In particular, a previously unnoticed cluster of rapidly and transiently PAMP-induced genes is enriched in transcription factors whose functions, previously associated with biological processes as diverse as abiotic stress adaptation and stem cell activity, appear to converge on growth restriction. Furthermore, some examples of known potentiators of PTI, in one case under direct MAP kinase control, support the notion that the rapidly induced transcription factors could constitute direct links to PTI signaling pathways and drive gene expression changes underlying establishment of the immune state.

## INTRODUCTION

Pathogen-associated molecular patterns (PAMPs) are conserved molecules or molecular assemblies that satisfy two criteria: (i) they are required for essential cellular or physiological functions of a pathogen and are therefore bound to evolve slowly, and (ii) they do not exist in the hosts of the pathogen (Janeway 1989; Medzhitov and Janeway 1997). Therefore, PAMPs constitute targets for distinguishing non-self from self by host immune receptors. Such PAMP-triggered immunity (PTI) mediated by specific receptors is crucial for pathogen defense in plants and animals (Nürnberger et al. 2004). In plants, a limited number of PAMP receptors have been identified that recognize conserved bacterial or fungal structures, such as flagellin (Felix et al. 1999), Elongation Factor Tu (EF-Tu) (Kunze et al. 2004; Zipfel et al. 2006), lipopolysaccharide (LPS), and chitin (Kaku et al. 2006; Miya et al. 2007), or even pathogen-induced aberrant host molecules such as oligosaccharides released from fungal digestion of the plant cell wall (D’Ovidio et al. 2004). The PAMP receptors FLAGELLIN-INSENSITIVE2 (FLS2) and EF-Tu RECEPTOR (EFR) recognize conserved peptides in bacterial flagellin (flg22) (Gómez-Gómez and Boller 2000; Chinchilla et al. 2006) and Elongation Factor Tu (elf18) (Kunze et al. 2004; Zipfel et al. 2006), respectively. These receptors contain an extracellular ligand-binding domain with leucine-rich repeats (LRRs) and a cytoplasmic serine/threonine kinase domain (Gómez-Gómez and Boller 2000; Chinchilla et al. 2006); (Kunze et al. 2004; Zipfel et al. 2006)). Binding of the ligand initiates a signaling pathway that implicates a host of co-receptors, and involves mitogen-activated protein (MAP) kinase cascades (Asai et al. 2002; Chinchilla et al. 2007). Important MAP kinase substrates include transcription factors (TFs) of the WRKY class (Andreasson et al. 2005; Qiu et al. 2008; Mao et al. 2011), so named after their invariant Trp-Arg-Lys-Tyr tetrapeptide implicated in DNA binding (Eulgem and Somssich 2007). WRKY transcription factors are themselves early PAMP-response genes that regulate many defense genes characterized by the presence of WRKY-binding sites (W-boxes) in their promoters (Eulgem and Somssich 2007). It has also recently been shown that PTI activation in *Arabidopsis* involves MAP kinase-dependent alternative splicing (Bazin et al. 2020) and translational reprogramming dependent on several features, including occurrence of a specific sequence element in 5’-leaders and sometimes skipping of upstream open reading frames (uORFs) in mRNAs encoding immune regulators (G. Xu et al. 2017; Pajerowska-Mukhtar et al. 2012).

Despite extensive gene expression profiling studies of plant PTI, a number of fundamental questions regarding the precise nature of transcriptional reprogramming and its control remain unresolved. These questions include, but are not limited to, three distinct areas. First, it remains ill-defined how early signaling events, such as MAP kinase activation that follows within less than five minutes of PAMP perception (Mészáros et al. 2006), are molecularly linked to transcriptional reprogramming that is typically measured, at the earliest, 30 minutes after PAMP perception (de Torres et al. 2003; Zipfel et al. 2004; Navarro et al. 2004; Ramonell et al. 2005; Truman, de Zabala, and Grant 2006; Zipfel et al. 2006; Moscatiello et al. 2006). Specifically, the time gap between MAP kinase activation and documented transcriptional responses suggests that earlier changes in gene expression may be part of the activation of PTI. Second, although it is now clear that alternative use of transcription start sites (TSS) has important ramifications for gene function in plant biology (Ushijima et al. 2017; Kurihara et al. 2018), no information regarding the extent of alternative TSS usage following PTI activation is available. Third, it is unclear whether immunity-related enhancers exist and are used to orchestrate the transcriptional activation of regulators and downstream response genes in Arabidopsis, as has been observed during activation of animal innate immunity (Arner et al. 2015). Answers to those questions should be tangible by gene expression profiling using Cap Analysis of Gene Expression (CAGE). In addition to information on transcript abundance, CAGE yields TSS information at nucleotide resolution, because it involves capture of capped transcripts, and generation of ~30 bp sequence reads immediately 3’ to the capped nucleotide (H. Takahashi et al. 2012). Thus, CAGE-based gene expression profiling clearly has the potential to answer questions on the possible existence of a very early PAMP-induced gene set, and on the possible use of PAMP-triggered alternative transcription initiation. Perhaps less intuitively clear is the fact that CAGE also has the potential to answer the third question on the possible existence of the PAMP-triggered enhancers. This is because of recent findings that enhancers in animal cells can be identified as DNAse Hypersensitive Sites (DHSs) that produce so-called enhancer RNAs (eRNAs): Bidirectional, short-lived transcripts, best observed upon inactivation of the RNA exosome complex (Andersson et al. 2014; Andersson and Sandelin 2020) which is at the core of cellular RNA processing and degradation by 3’-5’-exonucleolysis (Chlebowski et al. 2013). Because the study of enhancers in plants is still rudimentary, it is not yet clear whether eRNA-like-producing loci correspond to active enhancers. Nonetheless, intergenic and intronic loci producing eRNA-like transcripts have been observed in cassava using nascent RNA sequencing techniques (Lozano et al., n.d.), and we previously identified around 100 similar loci in unchallenged *Arabidopsis* seedlings using CAGE (Thieffry et al. 2020). In this latter case, detection of the short-lived eRNAs required inactivation of components of the nuclear exosome-mediated RNA decay pathway, achieved either by knockout mutation of the DEAD box helicase HUA ENHANCER2 (HEN2), a requisite, nucleoplasmic exosome cofactor (Lange et al. 2014), or by partial loss of function of the core exosome subunit RRP4 (Hématy et al. 2016; Thieffry et al. 2020). Thus, a search for PTI-related enhancers by an eRNA-focused approach would require inactivation of the nuclear exosome-mediated RNA degradation. Incidentally, in addition to facilitation of eRNA detection, an assessment of the relevance of nuclear exosomal RNA decay for PTI activation has merit on its own. In yeast, there is evidence that mechanisms of alternative transcription termination coupled to exosome-mediated nuclear pre-mRNA decay contribute to shape rapid reprogramming of gene expression in response to stress (Bresson et al. 2017), and a recent study in Arabidopsis provided an example of requirement of HEN2 for expression of a specific intracellular immune receptor (RPS6) (Takagi et al. 2020), belonging to the Nucleotide Binding-Leucine Rich Repeat (NB-LRR)-class well known to be induced in PTI (Navarro et al. 2004).

In the present study, we conducted a time course CAGE profiling experiment in which wild type and *hen2* knockout mutants were analysed at 10 minutes after FLS2 activation by flg22, in addition to 30 minutes at which time the PTI response is known to be activated (Navarro et al. 2004). Although our efforts neither revealed extensive immune-responsive enhancer transcription, nor a substantial implication of nuclear exosomal RNA decay in PTI-induced reprogramming of gene expression, two conclusions of broad significance were reached. First, TSS change is widespread in PTI-induced genes. This includes TSS changes of potential functional significance in regulatory genes and defense effectors. Second, PAMP-induced reprogramming of gene expression involves hitherto overlooked events: The rapid induction of a large fraction of the known PTI response much earlier than previously appreciated, including transient induction of a set of mRNAs encoding regulatory proteins enriched in transcription factors. Functions of many of these early response genes appear to converge on restriction of cellular growth and division. These findings add substantial new insight into PAMP-triggered transcriptional reprogramming, and provide a detailed basis for the design of future studies to gain a molecular understanding of how early PTI signaling events control transcriptional reprogramming leading to establishment of the immune state.

## RESULTS and DISCUSSION

### Validation of PTI Induction and TSS Identification by CAGE

To study the PAMP-triggered immune response, we transferred 14-day old sterile-grown seedlings to liquid culture and applied 3.3 μM of flg22 peptide after two days of acclimation to the liquid medium. In addition to wild type and *hen2-4* (Lange, Sement, and Gagliardi 2011), we included seedlings of two different genotypes: the flg22-insensitive *fls2* mutant for initial validation of our experimental set-up, and a hypomorphic mutant allele of the exosome core factor RRP4 (*rrp4-2*) for CAGE profiling at 30 minutes to answer the additional question of whether nuclear RNA quality control mediated by the exosome machinery may play a role in PAMP-triggered reprogramming of gene expression (**Figure 1A**). Preliminary controls showed that flg22 treatments were effective, because known response genes such as FRK1, MPK3, WRKY22, and WRKY29 were induced in wild type, while no induction could be detected in the *fls2* mutants (**Supplemental Figure 1A**). We therefore constructed and sequenced triplicate CAGE libraries from flg22 inductions conducted in this way. Analyses of the resulting data from unchallenged samples have been reported previously (Thieffry et al. 2020). The present full data set, including flg22 induction, was treated in the same way, i.e. the 5’-ends of CAGE tags located within 20 bp from each other on the same strand were clustered into CAGE tag clusters (TCs), quantified using their total number of tags, and finally normalized into tags per million (TPM) per sample (**Supplemental Dataset 1**). Initial analyses showed that our CAGE data faithfully captured known TSSs (Thieffry et al. 2020), and delivered two arguments that our flg22 treatments induced global gene expression changes typical of PTI (**Figure 1B-D**). First, multidimensional scaling (MDS) of CAGE TCs showed that the samples mainly clustered according to time after flg22 induction and according to their genetic background (**Figure 1B**). Second, differential expression analysis identified more than 2,000 up-regulated CAGE TCs (log_2_ fold-change ⩾ 1, *FDR* ⩽ 0.05, see Methods) when comparing samples harvested 30 minutes post-treatment to untreated controls (**Figure 1E**, **Supplemental Dataset 2**). The set of genes with up-regulated CAGE TCs at 30 minutes was highly enriched in biological processes related to stress and defense, with hallmarks of PTI (**Figure 1C**).

**Figure 1.**
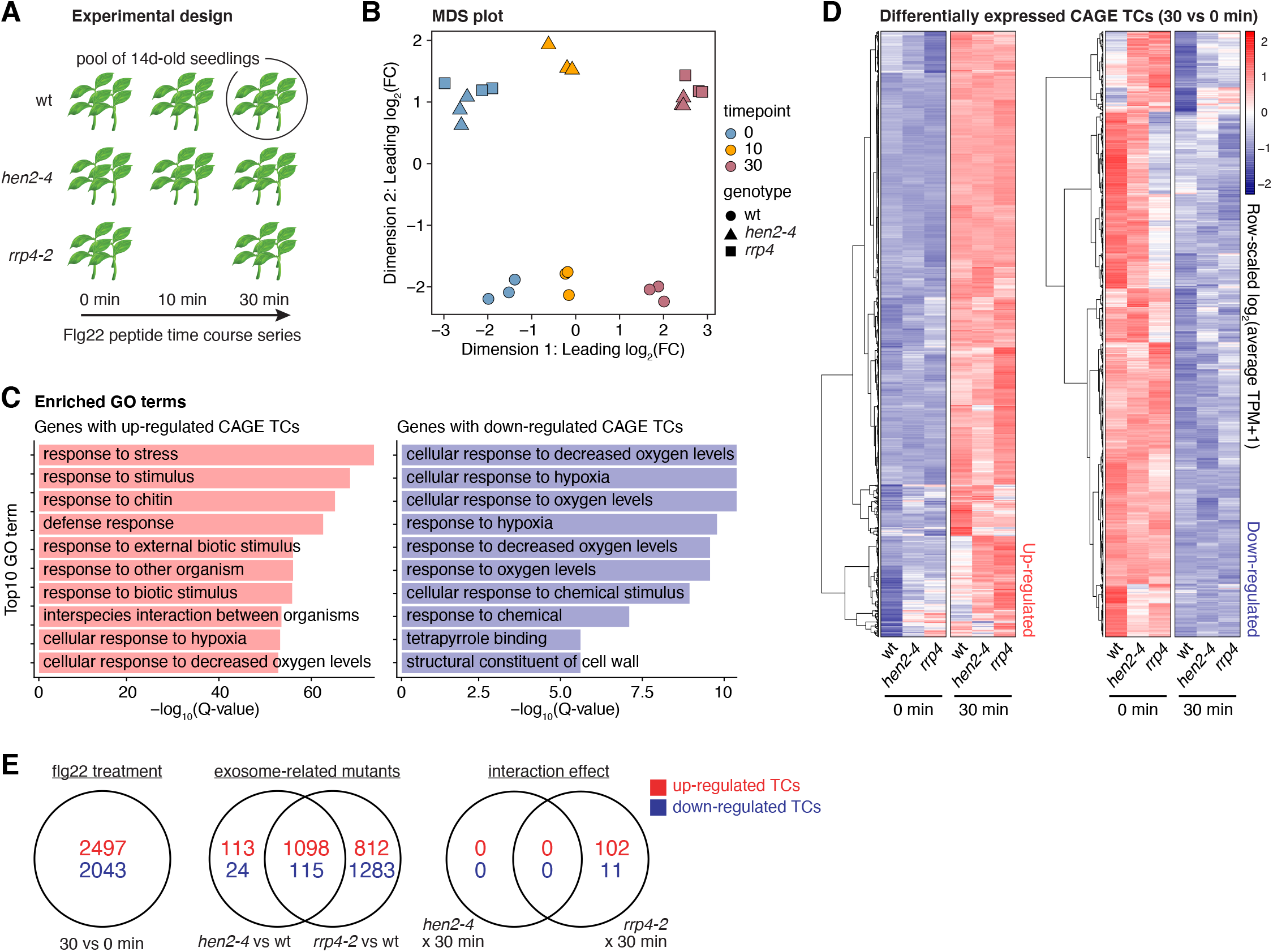
Experimental setup and validation of PTI response. **(A)** Overview of experimental design. Pools of 14d-old seedlings from a wild type (wt) and two exosome-related mutants (*hen2-4* and *rrp4-2)* were subjected to flg22 PAMP treatment in biological triplicates followed by CAGE library preparation and sequencing at set time points. Additionally, the *fls2* mutant was included in the time course for initial validation of the treatment. **(B)** Multi-dimensional scaling (MDS) plot of CAGE libraries. X and Y axes show the first two MDS dimensions. Each point corresponds to a CAGE library, colored by time of flg22 treatment. Shapes indicate genotype. Axes are scaled as leading log_2_ fold-change (FC); the root-mean-squared average of the log_2_ FC of the top 1000 genes best separating each sample. **(C)** Enriched gene ontology (GO) terms within genes with CAGE TCs responding to the flg22 treatment after 30 minutes. X-axis shows P-values after correction for multiple testing and −log base 10 transformed. Y-axis shows the top 10 enriched terms. Left and right panels show terms for genes with up-regulated (red) and down-regulated (blue) CAGE TCs. **(D)** Hierarchical clustering of CAGE tag clusters (TCs) induced at 30 minutes. CAGE TCs that were significantly up- or downregulated at 30 min after flg22 treatment vs 0 minutes were hierarchi-cally clustered. Left heatmap shows up-regulated CAGE TCs, right heatmap shows down-regulated. Rows show TCs. Columns show genotypes and time. Cell color indicates row-scaled TPM-normalized expression for respective TC in a given genotype and time point. **(E)** Venn diagram of differentially expressed CAGE TCs across experimental conditions. Red and blue numbers show up- and down-regulation, respectively. flg22 treatment shows TCs responding to the flg22 induction at 30 minutes compared to control (0 minute). The ‘exosome-related mutants’ denotes the comparisons of *hen2-4* and *rrp4-2* to wild type samples. The ‘interaction effect’ captures TCs whose response differs due to the interaction of exosome-related (*hen2-4* or *rrp4-2*) mutants and flg22 treatment at 30 min.

### Nuclear Exosome-mediated RNA Decay does Not Contribute Substantially to PTI-associated Reprogramming of Gene Expression

We next compared the PTI response in wild type with that in *hen2-4* and *rrp4-2*. We found that the overall PTI response was very similar in wild type and *hen2-4* (**Figure 1D**), including induction of MPK3, WRKY22 and WRKY29 as in wild type (**Supplemental Figure 1B**). Nonetheless, a small set of around 100 CAGE TCs was differentially expressed between wild type and *rrp4-2* at 30 minutes post-treatment (**Figure 1E**) (**Supplemental Dataset 2**). Since HEN2 is strictly nucleoplasmic, and RRP4 is a core exosome component required for both nuclear and cytoplasmic exosome functions, the most straight-forward interpretation of these results is that exosome-mediated cytoplasmic mRNA decay plays a minor role in induction of the immune state in *Arabidopsis*. In contrast, there is little indication for such a role of nuclear RNA quality control. These are relevant conclusions, because nuclear RNA quality control mediated by the exosome has previously been observed to have such regulatory roles in stress-induced genetic reprogramming in yeast (Bousquet-Antonelli, Presutti, and Tollervey 2000; Bresson et al. 2017), as well as transcriptional control in mouse stem cells (Lloret-Llinares et al. 2018; Garland et al. 2019). We also note that the possible involvement of cytoplasmic exosomal mRNA decay is consistent with recent identification of mammalian mRNAs whose degradation depends exclusively on the cytoplasmic exosome pathway (Tuck et al. 2020). It is also consistent with our previous identification of an extensive set of highly expressed mRNAs that overaccumulates relative to wild type in *rrp4-2*, but not in *hen2-4*, in untreated seedlings (Thieffry et al. 2020).

### Alternative TSS Usage During PAMP-triggered Genetic Reprogramming is Common

Although many cases of alternative transcription initiation may be ascribed to noisy start site definition by the RNA polymerase II pre-initiation complex (C. Xu, Park, and Zhang 2019), alternative promoter usage may increase the repertoire of gene regulation in a number of ways. First, multiple promoters resulting in the same protein product may facilitate distinct responses to different stimuli, or jointly increase the dynamic range of expression. Second, alternative promoters may be located so that the resulting transcripts contain different RNA regulatory elements or encode distinct sets of functional protein domains or localization signals. Recent studies show that this type of regulation has profound importance in the plant response to light or sensing of light quality (Ushijima et al. 2017; Kurihara et al. 2018). Nonetheless, the extent of alternative promoter usage in the PTI response is not known. To fill this gap, we first considered the pan-experiment landscape of alternative TSSs by investigating the localization of intragenic CAGE TCs regardless of expression dynamics. Similar to our previous analysis on untreated seedlings (Thieffry et al. 2020), a clear majority (90%) of detected genes had only one CAGE TC that contributed at least 10% to their host gene expression (this constraint was used for all intragenic TCs analyzed below, see Methods) (**Figure 2A**, **Supplemental Dataset 1**). Using a simplified hierarchical annotation strategy (**Figure 2B**, right), we observed that TCs in such single-TC genes were predominantly located within the promoter region (CAGE TC peak located ± 100 bp from TAIR10 annotated TSSs) of protein-coding and non-coding genes alike, as expected (**Figure 2B**). For example, ~90% of TCs in single-TC gene overlapped the promoter region and only ~6% were located within gene bodies. For genes with multiple TCs, we labelled the most highly expressed TC as ‘major’ (others as ‘minor’), and overlapped those with the simplified annotation as above. Both major and minor TCs were most commonly observed in annotated promoter regions, but a substantial fraction was also observed within gene bodies. For example, in protein-coding genes, major and minor TCs overlapped annotated promoter regions in 72 and 43% of cases, and gene bodies in 22 and 45% of cases, respectively (**Figure 2B**). These observations indicate that although the great majority of genes only use one TC, a considerable number of alternative TCs in multi-TC genes are found within gene bodies, motivating a more in-depth analysis of alternative TSS usage in PTI.

**Figure 2.**
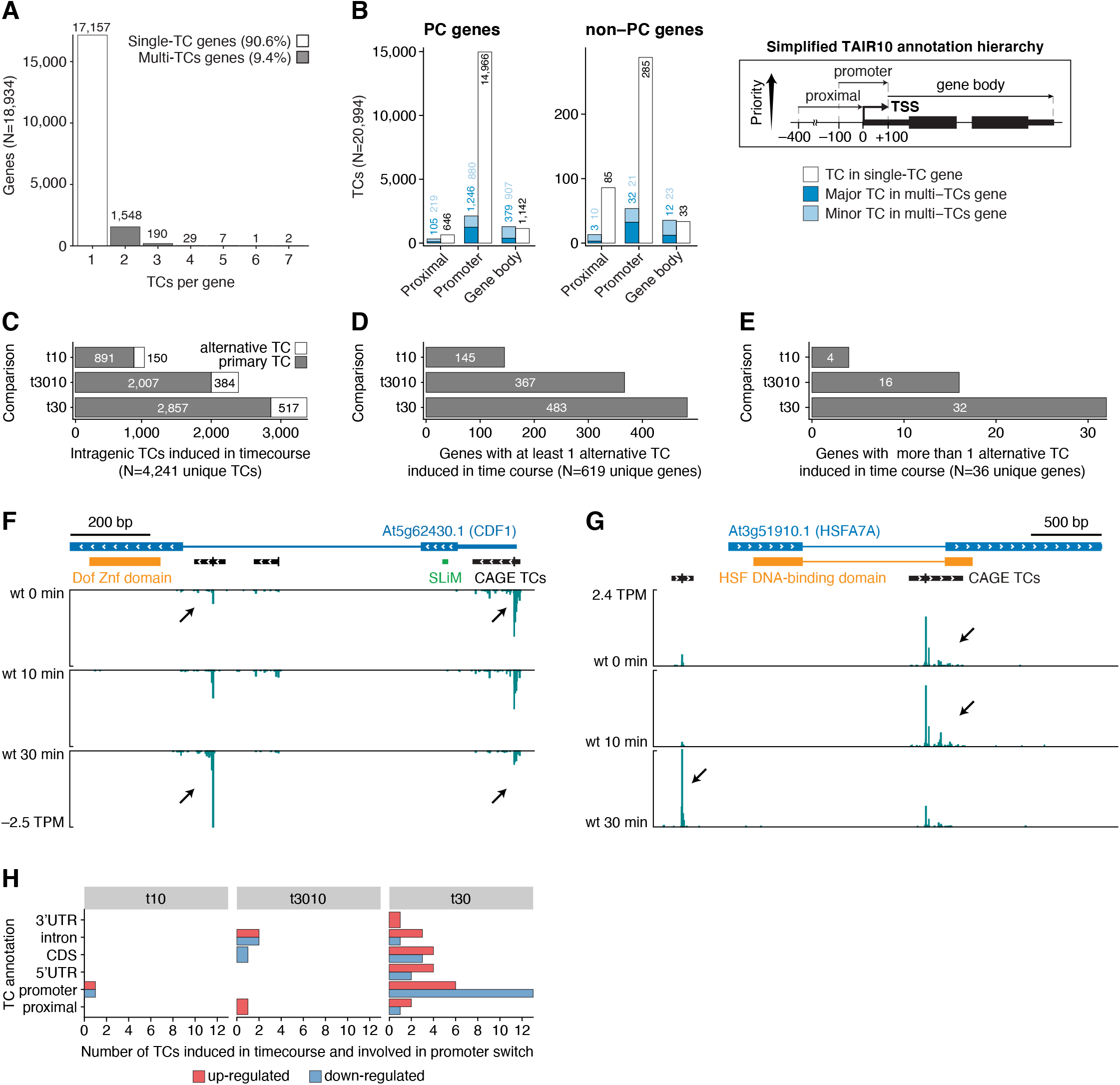
Alternative TSS usage in PTI. **(A)** Extent of alternative TSS usage. Y axis shows the number of genes having a set number of TCs (X-axis). Only intragenic TCs contributing at least 10% to their gene expression were considered (see Methods). Bar fills indicate whether genes have only a single TC (no fill) or multiple TCs (filled). **(B)** Annotation of intragenic TCs. Number of TCs (Y-axis) overlapping TAIR10 genomic features (X-axis) based on a simplified hierarchical annotation system (right). Left panels show data for protein-coding (PC) genes, middle panels for non-PC genes. Bar colors indicate whether the TC category originates from a single- (white) or multiple-TC gene (blue), and whether the TC is the major (dark blue) or a minor contributor (light blue) to its gene expression. **(C)** X-axis shows the number of intragenic CAGE TCs that are differentially expressed during the flg22 treatment time course. Y-axis shows the time point comparison used for differential expression analysis: t10 (10 vs 0 min), t3010 (30 vs 10 min), and t30 (30 vs 0 min). Bar colors indicate whether CAGE TCs are located within +/− 100 bp from the most upstream 5’-end of TAIR10 gene models (primary, grey) or not (alternative, white) **(D)** X-axis shows the number of genes with at least one alternative TC induced during the time course. Y-axis organized as in C. **(E)** X-axis shows the number of genes with more than one alternative TC induced during the time course. Y-axis organized as in C. (**F**, **G**) Genome browser views of *CDF1* and *HSFA7A* genes. The TAIR10 gene model is shown on top with large blue blocks indicating coding regions, and slim blue blocks indicating untranslated regions. Blue lines represent introns. White arrows indicate direction of transcription. Protein domains are shown in orange. CAGE TC are shown as black blocks with the tick marking the position of each TC peak. For *CDF1*, the short linear motif (SLiM) required for TOPLESS binding is indicated by a green block. Bottom tracks show the CAGE signal expressed in average tags per millions (TPM) across wt replicates for 0, 10, and 30 minutes following flg22 treatment. Positive and negative CAGE TPM values indicate positive and negative strand, respectively. Black arrows highlight important changes in CAGE TCs usage during the time course (see main text). **(H)** Annotation of CAGE TCs involved in promoter switches during flg22 time course. X-axis shows the number of TCs falling in each annotation category (Y-axis). Bar colors indicate down- (blue) or up-regulation (red) for each comparison in the time course (columns).

We found that more than 15% of intragenic TCs differentially expressed across treatment time points did not overlap the primary annotated TSS, defined as the most upstream promoter from TAIR10 (±100 bp). These TCs could be therefore considered as manifestations of changed alternative promoter activity during the time course (**Figure 2C**). A total of 619 genes had at least one such alternative TC that was differentially expressed between at least one pair of time points, where the 30 to 0 min comparison had the highest number of differentially expressed alternative TCs (**Figure 2D**). Interestingly, a small set of 32 genes had two or more alternative TCs differentially expressed in the comparison of samples treated for 30 min versus untreated samples (**Figure 2E**). We conclude that use of alternative TSSs is not an uncommon feature of the transcriptional PTI response and went on to analyze consequences of alternative TSS usage in more detail.

### Promoter Switching During PTI

A small set of 21 genes had pairs of TCs which were differentially expressed in opposite directions over time - in other words, TSS switching (**Supplemental Dataset 3**). The genes *CDF1* (At5g62430) encoding the transcription factor CYCLING DOF FACTOR1, and *HSFA7A* (At3g51910) encoding the HEAT SHOCK TRANSCRIPTION FACTOR A7A provide compelling examples of such TSS switches likely to have functional impact. In *CDF1*, the switch from an upstream to a downstream TSS during PTI induction favors production of an mRNA encoding a protein lacking at least an N-terminal short linear motif (SLiM, IKLFG) required for interaction with the transcriptional co-repressor TOPLESS (Goralogia et al. 2017) (**Figure 2F**), or potentially even some of the DNA-binding Zinc-finger depending on the exact site of translation initiation on the induced mRNA. This TSS switch is therefore predicted to change CDF1 function profoundly during PTI activation, either by modulating important protein-protein interactions or by flat out abrogation of transcription factor function. Conversely, transcription at *HSFA7A* switches from a TSS in the first intron to a TSS slightly upstream of the annotated promoter region (**Figure 2G**). The HSF DNA-binding domain of HSFA7A is encoded mostly in the first exon, suggesting that in this case, PTI-induced TSS switching results in preferential production of a functional, DNA-binding transcription factor and, potentially, repression of an interfering product with oligomerization, but not DNA binding properties (Guo et al. 2016). The promoter switching at the *HSFA7A* locus may contribute to PTI, because the HSF family plays important roles in plant stress adaptation (Guo et al. 2016). For example, HSFB1 is required for PTI induced by elf18, but not by flg22 (Pajerowska-Mukhtar et al. 2012). Overall, TSS switching events, often from a down-regulated TC in the promoter region to an up-regulated TC localized downstream, were mostly observed after 30 minutes of flg22 treatment (**Figure 2H**), perhaps suggesting that TSS switching may play a role in maintenance of the immune state rather than its establishment.

### Alternative TSS Usage Affecting Domain Composition

Inspired by the examples of promoter switching above, we conducted an extended analysis of TCs whose localization would predict disruption or exclusion of protein content. We found that 454 intragenic CAGE TCs in our data set were localized within or downstream of protein domains (in total, this corresponded to 428 genes). Of these, 127 TCs within 125 genes were differentially expressed during the time course, including *AFP1* (*ABI5 BINDING PROTEIN*, At1g69260) encoding a transcriptional co-repressor and *SUVR5* (At2g23740) encoding a histone H3 lysine 9 (H3K9) methyl transferase (**Figure 3 A, B**, **Supplemental Dataset 4**). AFP1 and SUVR5 have both been implicated as repressors of gene expression with importance in environmental adaptation. Mutation of *AFP1* causes abscisic acid hypersensitivity and reduced salt stress resistance (Garcia et al. 2008), while mutants in *SUVR5* display de-repressed expression of genes with gene ontology (GO) term “*Response to stimulus*”, including stress-response and auxin-response genes (Caro et al. 2012). Both *AFP1* and *SUVR5* show similar expression dynamics during PTI activation. A TC corresponding roughly to the annotated TSS that gives rise to an mRNA encoding full length protein is constitutively expressed, while a downstream TC is strongly repressed upon PTI activation (**Figure 3A, B**). In both cases, the downstream TSS directs production of an mRNA encoding a single protein domain that may interfere with the function of the full-length protein. In the case of AFP1, the putative truncated protein product contains only the transcription factor-binding Jas domain, but not the EAR (Ethylene-responsive binding factor-Associated Repression) and NINJA (Novel Interactor of JAZ) domains with transcriptional co-repressor function. Similarly, for SUVR5, the putative truncated protein contains only pre- and post-SET domains involved in substrate binding and regulation of catalysis, but neither of the Zinc finger domains required for DNA binding, nor the actual H3K9 methyltransferase domain (Caro et al. 2012).

**Figure 3.**
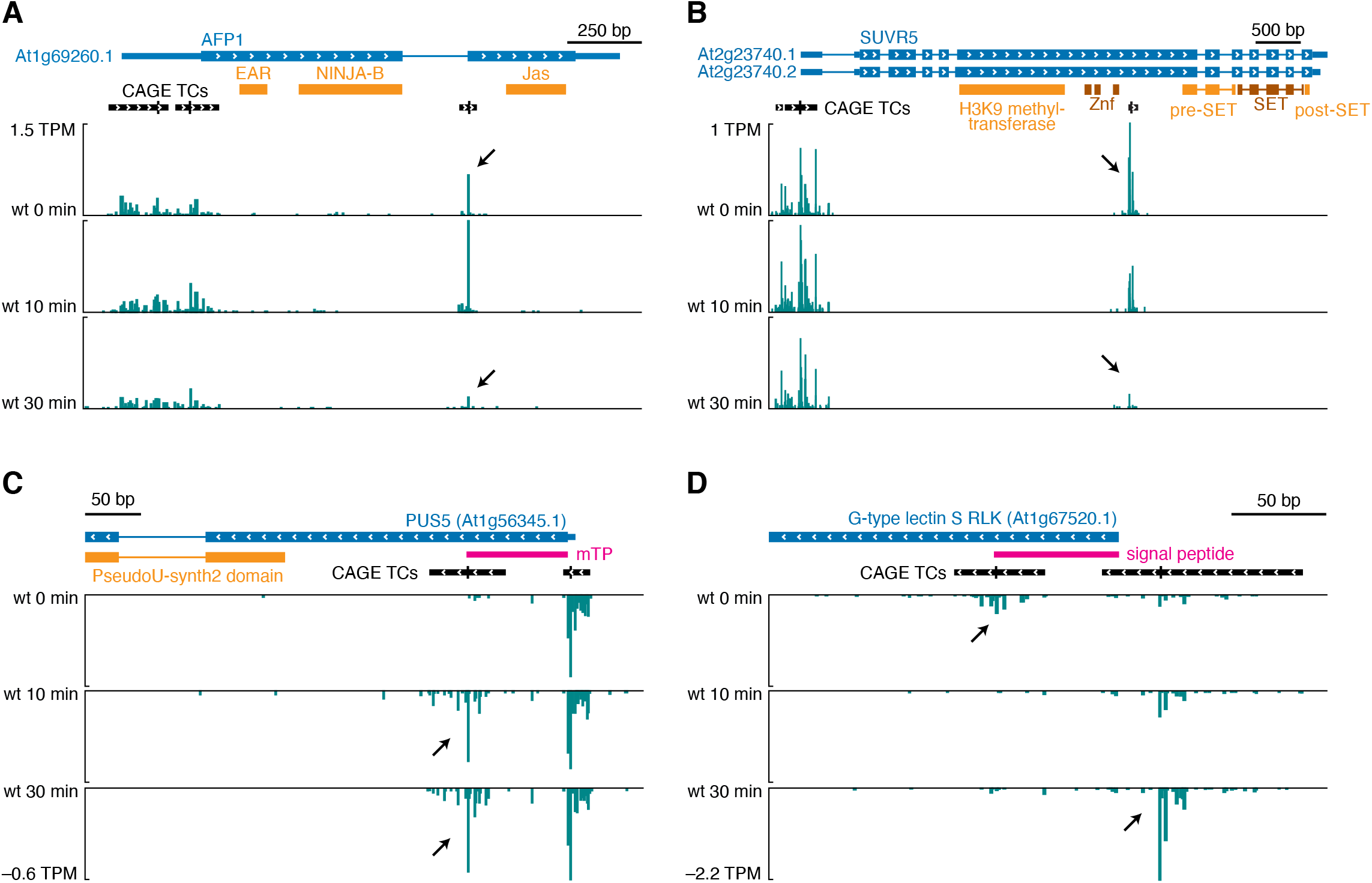
Examples of alternative TSSs excluding protein domains or influencing the presence of a target peptide. Genome browser views for *AFP1* (A), *SUVR5* (B), *PUS5* (C), and a G-type lectin S-RLK-encoding gene (D), organized as in **Figure 2F**. Pink blocks indicate predicted target peptide.

### Alternative TSS Usage Affecting N-terminal Target Peptides

To catalog transcription initiation events affecting the occurrence of predicted target peptides for entry into the secretory pathway, or for plastidial or mitochondrial import, we first scanned the complete set of TAIR10 proteins with SignalP-5.0 and TargetP-2.0 (Jose Juan Almagro Armenteros et al. 2019) (see Methods) and identified 6,985 genes with a predicted localization signal (**Supplemental Dataset 5**). Among those, 4% (284/6,985) had at least one intragenic CAGE TC (subjected to the same thresholds as above) located within or downstream of the sequence encoding a predicted target peptide. Notably, 78 of those TCs (within 76 genes) were differentially expressed in the flg22 time course (**Supplemental Dataset 5**). To further analyze the occurrence of differential exclusion of N-terminal target peptides, we identified intragenic TCs whose response in the time course differs from that of other TCs within the same gene (using limma:diffSplice, see Methods). Clear cases of such differential TC usage (DTU) involving partial or total exclusion of a localization signal were identified in 26 genes, where one promoter produced a transcript with and another without the signal peptides (**Supplemental Dataset 5**).

We consider two major possible consequences of differential TC usage around sequences encoding target peptides: the first is of potential regulatory importance, the other probably corresponds to molecular error correction. The first outcome is illustrated with the *PUS5* gene (At1g56345) encoding a stand-alone (i.e., snoRNA-independent) pseudouridine synthase, showcasing a downstream TC rapidly induced by flg22 (**Figure 3C**). In contrast to the constitutively expressed, annotated TSS, the mRNA resulting from transcription initiation at the induced downstream TC does not include the sequence encoding a mitochondrial transit peptide (**Figure 3C**). Importantly, it does retain coding potential for the pseudouridine synthase catalytic domain, thus raising the possibility that PUS5 performs cytoplasmic functions during PTI activation. Stand-alone pseudouridine synthases catalyze uridine-pseudouridine isomerization, an RNA modification whose occurrence is well established in tRNAs, but has also recently been found in mRNAs (Borchardt, Martinez, and Gilbert 2020) in several eukaryotic organisms, including plants (Sun et al. 2019). Given (i) the importance of rapid translational reprogramming for PTI (Pajerowska-Mukhtar et al. 2012; G. Xu et al. 2017), (ii) the relevance of tRNA modifications for translational control (Tuorto and Lyko 2016; Borchardt, Martinez, and Gilbert 2020), and (iii) the rapid nature of *PUS5* alternative TSS induction by flg22 (**Figure 3C**), it is an attractive possibility that alternative TSS usage at the *PUS5* gene contributes to translational reprogramming during PTI activation by directing cytoplasmic PUS5 function.

The second case involves a gene encoding an orphan G-type lectin S-receptor-like kinase (At1g67520). In the unchallenged state, this gene contains lowly expressed TCs both upstream and downstream of a signal peptide-encoding sequence (**Figure 3D**). In contrast, upon PTI activation, only one upstream and much more highly expressed TSS predominates, presumably giving rise to a functional receptor that enters the secretory pathway as it should (**Figure 3D**). This pattern of re-focusing low-level noisy transcription initiation, potentially giving rise to mRNA species encoding faulty protein, to a single, highly expressed TSS upon induction is probably an example of the recent proposition that alternative transcription initiation is often non-adaptive, and rather a consequence of imperfect TSS selection by the transcription initiation machinery (C. Xu, Park, and Zhang 2019).

### TSS Change Resulting in Transcripts Differing in uORF Content

Translational control is emerging as a major theme in regulation of plant stress responses (G. Xu et al. 2017; Kurihara et al. 2018). Transcript quality is important for the efficiency of mRNA translation, and recent studies have indicated that uORFs are implicated in control of gene expression during the activation of PTI as well as other stress responses (G. Xu et al. 2017). Since alternative TSS usage can lead to inclusion or exclusion of uORFs in mRNAs (Kurihara et al. 2018), we specifically searched for cases that would lead to production of such alternative mRNAs upon flg22 induction. Using the same approach as above, we found 510 TCs in 480 genes that were within or downstream uORF regions and differentially expressed during the flg22 time course (**Supplemental Dataset 6**). For example, chaperone induction is associated with PTI (Navarro et al. 2004), and our analyses show that during PTI activation, the genes encoding both an Hsp70 isoform (At1g16030, Hsp70.6) and an Hsp70 nucleotide exchange factor (BCL2-associated athanogene 6, BAG6, At2g46240) use downstream TSSs excluding one and three uORFs in their mRNAs, respectively (**Figure 4A, B**). Another illustrative example is provided by the gene encoding the GATA-type transcription factor BME3/GATA8 (At3g54810) (**Figure 4C**). In unchallenged wild type, this gene produces different alternatively spliced mRNAs with long 5’-leaders, one of which contains two uORFs. In contrast, upon PTI activation, transcription initiates downstream of the first intron and gives rise to an mRNA with a dramatically shortened leader free of uORFs, potentially leading to enhanced efficiency of translation. Finally, we note that in the gene encoding the important signal transducer MEKK1 (At4g08500), a MAP kinase kinase kinase required for activation of the MPK4 cascade by flg22 (Ichimura et al. 2006; Suarez-Rodriguez et al. 2007), the TSS distribution within its sole CAGE TC shifts during PTI activation so that a mRNA free of uORFs becomes predominant (**Figure 4D**). Taken together, our observations show that alternative use of TSSs is widespread in PTI induction and provide a solid basis for future functional studies on the consequences of alternative TSS usage for genes participating in regulation or execution of the immune state.

**Figure 4.**
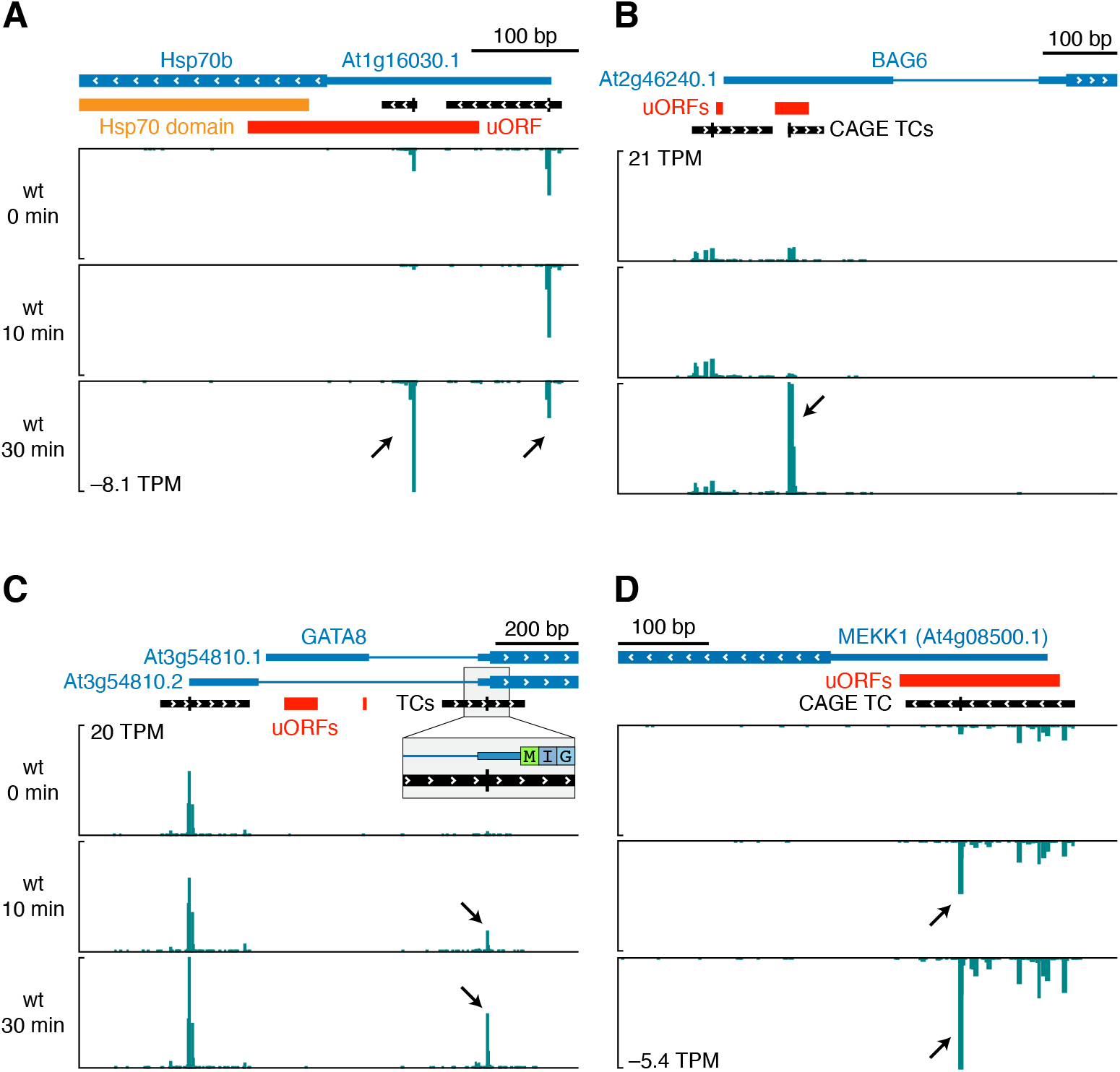
Examples of alternative TSSs influencing the presence of uORFs. Genome browser views for *Hsp70b* (A), *BAG6* (B), *GATA8* (C), and *MEKK1* (D) genes, organized as in Figure 2F. Red blocks indicate upstream open reading frames (uORFs).

### Activation of eRNA-like Transcription is Not Widespread in PTI induction

We next addressed the question of possible existence of PTI-activated enhancers revealed by eRNA transcription. To identify such potential enhancers, we searched for short (~ 500 bp) loci featuring divergent transcription in at least 3 samples (see Methods). We found 155 enhancer candidates located in intergenic or intronic regions (**Supplemental Dataset 7**) of which 15 produced higher RNA levels in *rrp4-2* or *hen2-4* than in wild type (**Supplemental Figure 2A, B**, **Supplemental Dataset 2**, see Methods). However, only four bidirectionally transcribed loci were induced upon flg22 treatment (**Supplemental Figure 2B**), and only a single locus showed the behaviour expected from putative PTI-activated enhancers, i.e. sensitivity to exosome mutation and induction by flg22 (**Supplemental Figure 2B**). We conclude that our eRNA-focused approach did not reveal widespread existence of PTI-related enhancers.

### The Known PTI Response is Preceded by Rapid and Transient Induction of Regulatory Genes

Finally, we explored the CAGE data for patterns of gene expression change over the time course. To this end, we aggregated CAGE TCs across gene models and conducted differential expression analysis with limma (log_2_ fold-change ⩾ 1, *FDR* ⩽ 0.05, see Methods) to identify genes responding during the flg22 treatment time course. Differentially expressed genes (DEGs) were then classified into induced or repressed sets based on hierarchical clustering of normalized gene expression values (**Figure 5A**, **Supplemental Dataset 8**). This analysis defined three clusters in each category (induced, clusters 1, 2, 5; repressed, clusters 3, 4, 6) according to their expression trajectory over time (**Figure 5B**). Cluster 1 (C1), characterized by genes activated only at 30 minutes after flg22 stimulation, contained the large set of previously characterized PAMP response genes, including WRKY transcription factors as well as defense effectors such as chitinases and other pathogenesis-related (PR) genes. Consistent with these molecular functions, the C1 gene set was enriched in the GO terms “*response to stress, biotic stimulus, and response to other organisms”* (**Figure 5C**, see Methods). Genes in cluster 2 (C2) responded already after 10 minutes and their expression continued to increase over the 30 minute time course. Thus, a sizable fraction of the known PAMP response is activated much earlier than appreciated until now, and is in temporal proximity to early signal transduction events such as MAP kinase activation. Intriguingly, cluster 5 (C5) genes were rapidly induced at 10 minutes, but had returned to basal expression levels at 30 minutes. The transient nature of C5 induction means that C5 genes may have been largely overlooked in previous profiling studies of the PAMP transcriptional response. Thus, C5 has particular potential to reveal new aspects of transcriptional reprogramming upon PAMP perception.

**Figure 5.**
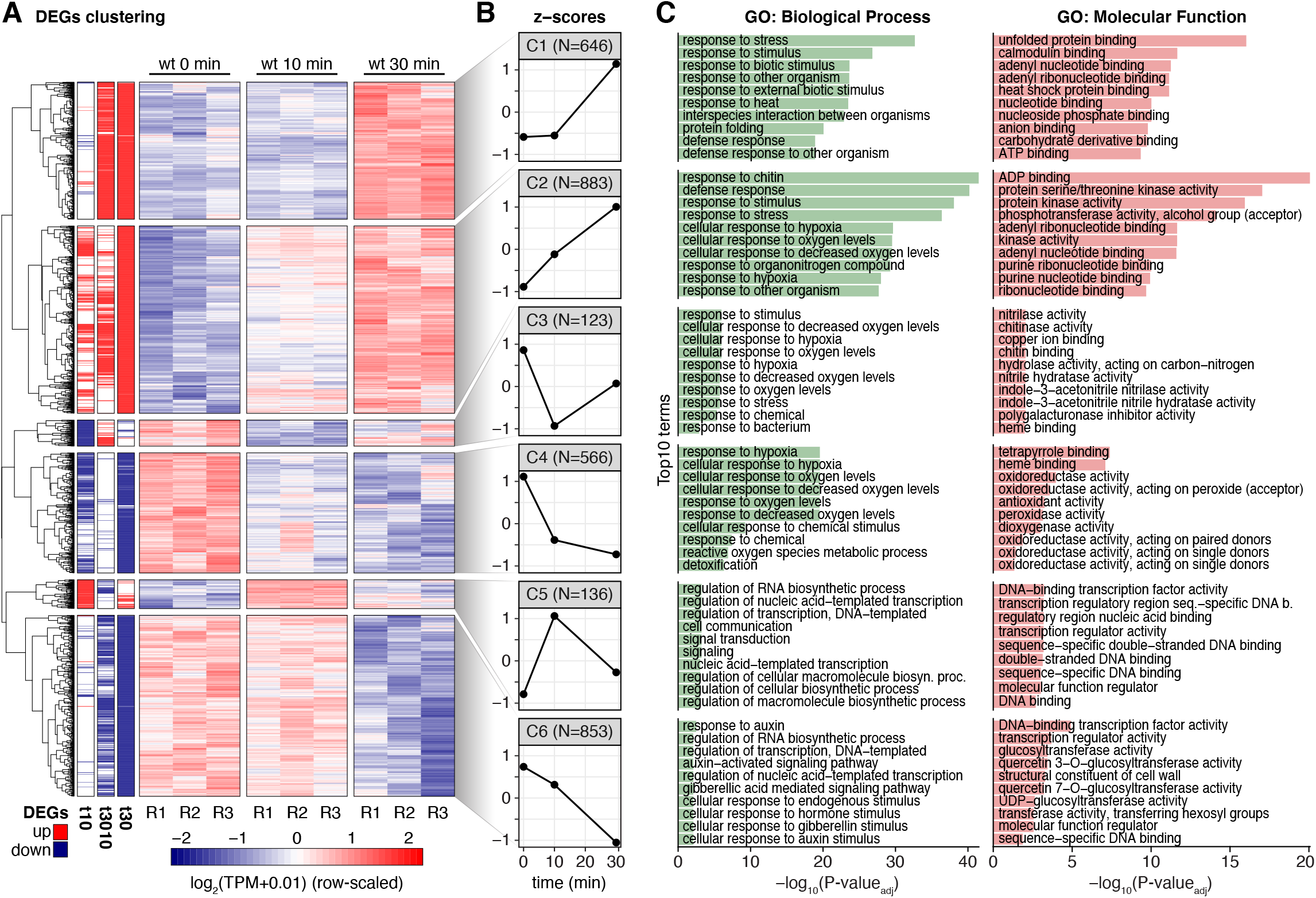
Expression clustering of the PTI transcriptional response. **(A)** Heatmap of differentially expressed genes (DEGs) in flg22 treatment. Each row represents a gene differentially expressed in the flg22 induction time course. Left three columns indicate up- (red) or down-regulation (blue) status according to the comparisons (30 vs 0 min, 30 vs 10 min, 10 vs 0 min). Remaining columns show replicates for each time point for wt samples. Colors represent normalized CAGE expression (log_2_TPM+0.01). Six expression patterns were identified by hierarchical clustering, numbered from top to bottom (C1-C6). **(B)** Average z-score of genes in clusters defined in A. X-axis shows treatment timepoints, in minutes. Y-axis shows the average expression z-score for each gene cluster from A. **(C)** Enriched gene ontology terms of genes in clusters defined in A. Organized as in Figure 1C, but both Biological Processes (left panel), and Molecular Function (right panel) GO term results are shown.

To further validate these central observations on the temporal nature of reprogramming of gene expression in PTI activation, we performed independent flg22 inductions and analyzed a slightly extended time course series (0, 10, 30, and 60 minutes after flg22 addition) by standard RNA-sequencing (RNA-seq). We found similar expression trends for the sets of genes in CAGE-defined clusters, albeit often with a temporal lag compared to the CAGE data (**Supplemental Figure 3**). The lag is probably explained by the fact that CAGE detects not only mature mRNAs but also pre-mRNA species, because it relies on 5’-cap capture in combination with random-primed reverse transcription. RNA-seq, on the other hand, uses oligo-dT-selection prior to reverse transcription so that (pre-)mRNAs must have undergone 3’-end formation to be detected. Taken together, our results reveal a hitherto unknown temporal order of reprogramming of gene expression during PTI activation. Many final PAMP response genes, including regulatory factors (**Supplemental Dataset 8**), are induced immediately after elicitation. Intriguingly, this immediate wave of gene induction also includes transiently induced genes (C5) that we analyze in more detail below due to its outstanding potential to reveal novel aspects of PTI activation.

### Cluster 5 is Enriched in Transcription Factors

Inspection of the functional annotation of genes contained in C5 showed that it was significantly enriched in GO terms related to transcription factors and nucleic acid binding activities (*FDR* ≤ 0.05, see Methods) (**Figure 5C**). Indeed, compared to other clusters of flg22 responsive genes, the fraction of genes encoding known transcription factors in C5 was roughly 3-fold higher (**Figure 6A**, see Methods). Because of the potential of the early-induced transcription factors to orchestrate the eventual transcriptional output in PTI, we focused our efforts on understanding the relevance of C5 transcription factors.

**Figure 6.**
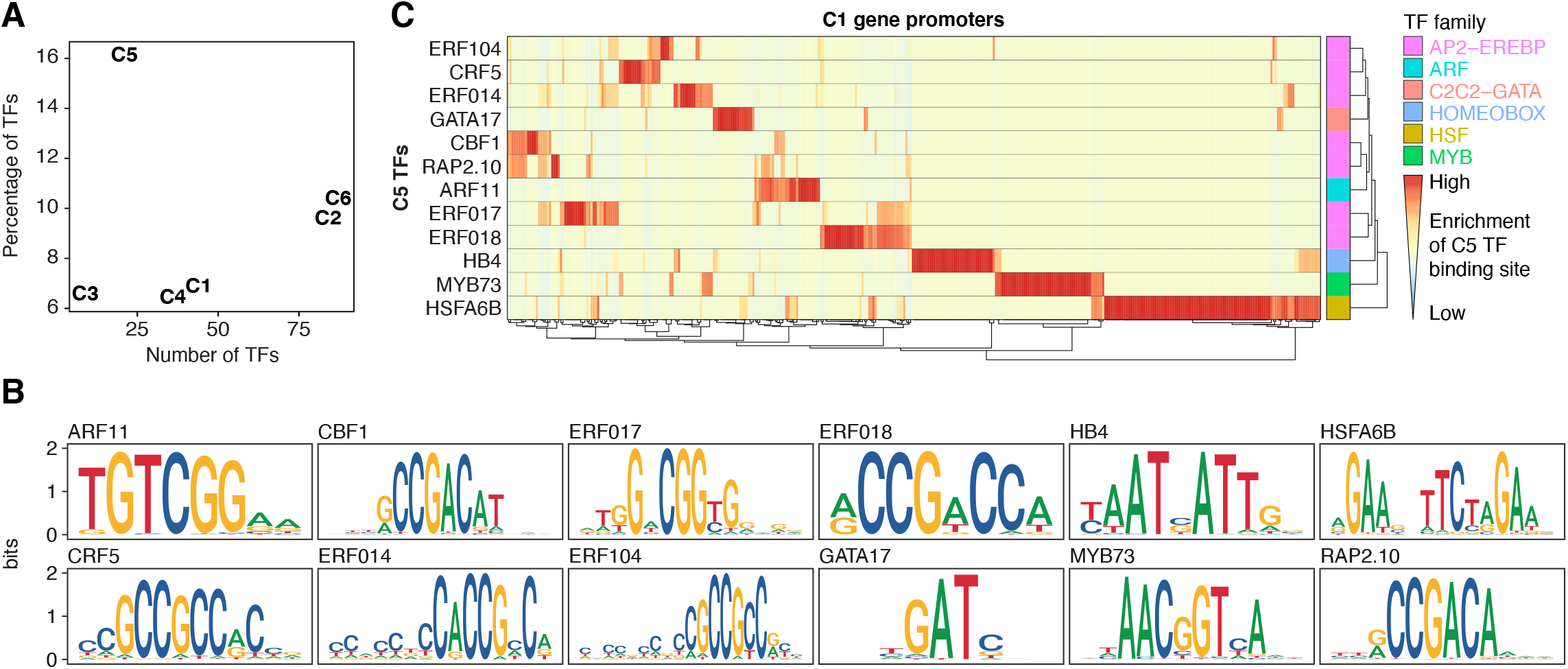
Promoters of PAMP response genes are enriched in C5 transcription factor binding sites. **(A)** Known transcription factor genes in DEG clusters (Figure 5A). X-axis shows the number of detected transcription factors (TFs) in each of the expression clusters from Figure 5A. Y-axis shows the proportion of TFs for each cluster. **(B)** Sequence logo plots of cluster 5 transcription factors for which a profile could be retrieved from the JASPAR database. Y-axis shows information content in bits. **(C)** Enrichment of binding sites from C5 transcription factors in the promoter regions of C1 genes. Heatmap rows indicate cluster 5 TF models, whereas columns represent C1 promoter regions. Heatmap colors show the ratio of binding site matches to the genome-wide average in all promoters (see Methods), with red indicating high over-representation. Right-most column shows the family of respective TF. Similar enrichment analysis for genes in all differentially expressed clusters is shown in **Supplemental Figure 4**.

### Links of C5 Transcription Factors to PTI Signaling and Establishment of Immunity

We first noted that several C5 transcription factors have been implicated in PTI (ETHYLENE RESPONSE FACTOR104 (ERF104, At5g61600, (Bethke et al. 2009); ERF014 (At1g44830, (H. Zhang et al. 2016); MYB34 (At5g60890, (Frerigmann and Gigolashvili 2014; Frerigmann et al. 2016)) or other branches of plant immune responses to fungal and bacterial pathogens (ERF016 (At5g21960) (Zhao et al. 2021)). Strikingly, the ERF104 transcription factor provides an example of a direct link to a major flg22-activated signal transducer, the MAP kinase MPK6. The stable MPK6-ERF104 complex dissociates within 5-15 minutes of flg22 perception (Bethke et al. 2009), closely matching the kinetics of MPK6 activation (Mészáros et al. 2006) and transcriptional activation of *ERF104* observed here. In addition, both knockout mutants and overexpressors of *ERF104* exhibit enhanced bacterial susceptibility (Bethke et al. 2009), pointing to the physiological relevance of the transient induction we describe here. It is also noteworthy that knockdown of ERF014 delays flg22-responsive gene expression while its overexpression is sufficient to cause bacterial resistance and hyperresponsiveness to flg22 perception (H. Zhang et al. 2016). These observations support the notion that an important function of at least some C5 transcription factors is to directly link PTI signal transducers to transcriptional reprogramming and potentiation of the immune state.

### C5 Contains Regulators of General Stress and Stem Cell Properties

ERF104 is also induced within minutes in response to abiotic stresses that require growth arrest (Moore, Vogel, and Dietz 2014; Vogel et al. 2014; Illgen et al. 2020), suggesting that it functions more generally in rapid stress adaptation than specifically in PTI activation. Indeed, despite their association with seemingly distinct biological processes (abiotic stress, biotic stress, stem cell functions), a common denominator of functions of many C5 transcription factors may be growth restriction as a general response to stress. Clear examples of this include at least five cases: (i) the key cold stress adaptation factors CBF1 (At4g25490) and CBF2 (At4g25470) (Liu et al. 2019), also shown recently to be induced by bacterial infection (Tuang et al. 2020), (ii) ERF017 (At1g19210) and ERF104, induced among other upon growth arrest-inducing intense light treatment (Vogel et al. 2014), (iii) ERF018/ORA47 (At1g74930) that has direct roles in control of biosynthesis of the growth-restricting phytohormones abscisic and jasmonic acid, and whose overexpression causes slow growth (Chen et al. 2016), (iv) ANAC044 (At3g01600), important for arrest of cell division in response to DNA damage (N. Takahashi et al. 2019), and (v) heat shock transcription factors HSFA3 (At5g03720) and HSFA6B (At3g22830) implicated in growth restriction and induction of chaperones destined to both cytoplasm and secretory pathways (Schöffl, Prändl, and Reindl 1998; Guo et al. 2016). In this regard, we note that the uORF-skipping alternative TSSs induced in Hsp70 and the nucleotide exchange factor BAG6 (**Figure 4**) may be linked to the immediate induction of HSFs in C5 and C2 (HSFA4A (At4g18880), HSFB2A (At5g62020), **Supplemental Dataset 8**), and heat shock transcription factors as a group have previously been proposed to be major drivers of the growth-to-defence transition (Pajerowska-Mukhtar et al. 2012).

The vascular stem cell-specific transcription factor WUSCHEL-like homeobox 4 (WOX4, At1g46480) (Etchells et al. 2013; Suer et al. 2011) may also belong to the category of growth-arresting factors, although it has not previously been associated with immunity or other stress responses. This is because of the recent finding that in stem cells of the shoot apical meristem, the founding WOX family member WUSCHEL (WUS) acts to repress translation globally via repression of ribosomal rRNA processing (Wu et al. 2020). Another stem cell transcription factor, KNAT1/BP (At4g08150) was also part of C5, although it is unclear whether its activities are related to repression of growth. Nonetheless, the combined induction of KNAT1 and WOX4 is intriguing for two reasons. First, KNAT1 and WOX4 act redundantly to control vascular stem cell activities (J. Zhang et al. 2019). Second, upon auxin stimulation of root pericycle cells, that exact transcription factor combination promotes establishment of a protective suberized, periderm layer rather than initiation of a proliferative stem cell niche destined to form a new lateral root (Xiao et al. 2020). Transcription factors associated with growth arrest and general stress responses were also found in C2 (e.g. HSFA4A, HSFB2A, CAMTA3 (At2g22300), CAMTA6 (At3g16940), ANAC062 (At3g49530), MYB74 (At4g05100), and several ERFs (Doherty et al. 2009; Vogel et al. 2014; Yang et al. 2014; Moore, Vogel, and Dietz 2014; R. Xu et al. 2015; Guo et al. 2016; Jacob et al. 2018; Illgen et al. 2020)), underscoring the multi-faceted reprogramming from growth and division to arrest immediately upon PAMP perception. Several additional transcription factors such as the Zinc finger/Homeobox factors HB4 (At2g44910) and HB28 (At3g50890) and the cytokinin response factor CRF5 (At2g46310, (Rashotte et al. 2006)) have not previously been associated with defence responses, and their inclusion in C5 therefore opens new venues to investigate their biological functions.

### Promoters of PAMP Response Genes are Enriched in Binding Sites for Cluster 5 Transcription Factors

We next asked whether C5 transcription factors other than the experimentally verified examples discussed above (ERF104, ERF014 and MYB34) had the potential to cause transcriptional reprogramming in PTI. If this were the case, promoters of PAMP response genes should show enrichment of C5 transcription factor binding sites. We retrieved position-specific weight matrix models describing DNA binding preferences of 12 transcription factors belonging to C5 from JASPAR CORE Plantae (Fornes et al. 2020) (**Figure 6B**, **Supplemental Dataset 9**, see Methods). Using these models, we computed the enrichment compared to the genome-wide average of predicted binding sites in the promoter regions of the genes of each of clusters 1-6 defined by differential gene expression in the flg22 time course (**Figure 5A**, see Methods). Interestingly, many promoters of genes activated or repressed in PTI were indeed enriched for predicted binding sites of C5 transcription factors (see **Figure 6C** for the known PAMP response genes (C1); **Supplemental Figure 4** for C1-C6). This also included C5 itself, perhaps suggesting that negative autoregulation contributes to the transient nature of its induction by flg22. In many cases, enrichment of elements corresponding to more than one C5 transcription factor could be identified, suggesting either a degree of functional redundancy between, or combinatorial binding of, C5 transcription factors in PTI. Nonetheless, we also identified many cases where only a single type of *cis*-element showed strong enrichment (**Figure 6C**). This pattern does not rule out overlapping functions of distinct transcription factors in regulation of PTI response genes, especially given the fact that many additional transcription factors (in our C2) are also induced long before the establishment of the immune state. Such a redundant setting of the system would be reminiscent of recent analyses of the transcriptional response to jasmonate in which inactivation of multiple early-induced transcription factors was required to observe measurable effects on the expression of later response genes (Hickman et al. 2017).

## CONCLUDING REMARKS

Our study provides substantial new insight into use of alternative transcription initiation sites and overall gene expression changes that take place rapidly after PAMP perception. The results should facilitate a better understanding of genetic reprogramming underlying the defence transition and establishment of the immune state in at least three ways: First, the discovery of very early PAMP response genes facilitates the design of studies aimed at linking immediate signal transduction events such as protein kinase activation directly to changed transcriptional output. It is possible that a refined temporal study to identify genuinely immediate responders, potentially through the use of nascent RNA techniques such as native elongating transcript sequencing NET-seq (Mayer and Churchman 2016; Zhu et al. 2018; Kindgren, Ivanov, and Marquardt 2020), combined with existing and refined knowledge on phosphoproteome changes following PAMP perception (Rayapuram et al. 2018), will be of value in this regard. Second, the crucial, but daunting task of deciphering key elements of the texture of the transcription factor web driving genetic reprogramming in PTI through genome-wide identification of binding sites is now tangible, because our results allow focus on a more limited number of early-responding transcription factors. Third, our study adds support to the importance of translational control in the early PAMP response (G. Xu et al. 2017; Pajerowska-Mukhtar et al. 2012), and hints that covalent modification of tRNAs and/or mRNAs could play roles in this regard.

## METHODS

### Plant Materials

All *Arabidopsis thaliana* plants are of the Col-0 ecotype. The *hen2-4* (At2g06990, SALK_091606C) mutant was described in (Lange, Sement, and Gagliardi 2011) and seeds were obtained from Dominique Gagliardi. The *rrp4-2* mutant is described in (Hématy et al. 2016) and was kindly provided by the authors. The *fls2* (At5g46330) mutant (T-DNA insertion line SALK_062054) was obtained from the Nottingham Arabidopsis Stock Centre (NASC).

### Genotyping

As in (Thieffry et al. 2020), DNA was isolated by adding one volume of phenol-chloroform (50:50 [v/v]) to freshly ground leaves in urea buffer (42% [w/v] urea, 312.5 mM of NaCl, 50 mM of Tris-HCl at pH 8, 20 mM of EDTA, and 1% [w/v] ɴ-lauroylsarcosine). Phases were separated and the DNA-containing supernatant was isolated. Nucleic acids were precipitated with isopropanol and pelleted DNA was rinsed with EtOH 70% (v/v). The DNA was used as a PCR template to confirm T-DNA insertion in *hen2-4* and *fls2* mutants. The single-point mutation in *rrp4-2* was confirmed by target DNA amplification and enzymatic digestion (Eco47I, *AvaII*). Genotyping primers are available in **Supplemental Dataset 10**.

### Growth Conditions

All growth conditions were as described in (Thieffry et al. 2020). Briefly, seeds were sterilized with 70% (v/v) EtOH, followed by 1.5% (w/v) sodium hypochlorite and 0.05% (w/v) Tween-20 (10 min, Sigma-Aldrich), then rinsed with sterile ddH_2_O. Clean seeds were stratified in complete darkness at 4°C for 72 hours, then germinated on 1% (w/v) Murashige & Skoog medium (supplemented with 1% [w/v] Suc and 0.8% [w/v] agar) under sterile conditions (Petri-dishes) with long-days light cycles (16-h L/8-h D photoperiod, 130 μmol photons m^−2^s^−1^ at 21◻, cat. No. Master TL-D 36W/840 bulbs; Philips). Intact 12 day-old seedlings were transferred in 8 mL of 1% liquid MS media (as above) in 6-well plates (Nunc, cat. No. 140675) and acclimated for 2 days with mild agitation (130 rpm), under identical light and temperature settings as above.

### Flagellin Treatments

Flg22 peptide with sequence Ac-QRLSTGSRINSAKDDAAGLQIA-OH was obtained from Schafer-N (www.schafer-n.com) with purity > 95%. Flg22 peptide was dissolved in dimethyl sulfoxide (DMSO) at 1 mg/mL. Biological replicates consisting of a pool of 10 seedlings were subjected to 3.3 μM of flg22 and 0.77% of DMSO under constant agitation (130 rpm) for 10 and 30 minutes for the CAGE samples, and 0, 10, 30, and 60 minutes for the RNA-seq samples. Seedlings were removed from the media and immediately flash-frozen in liquid nitrogen before undergoing RNA extraction.

### Total RNA Extractions

Total RNA was extracted as described in (Thieffry et al. 2020), with the addition of samples treated with flg22 peptide for 10, 30 and 60 minutes. Succinctly, plant materials were flash-frozen and TRI-Reagent (Sigma-Aldrich) was added to 100 mg of finely ground tissue. Following chloroform phase-separation, the aqueous phase was transferred to a fresh tube and the RNA was precipitated with one volume of isopropyl alcohol (400 μL) for 30 min at room temperature. Total RNA was pelleted by centrifugation (10 min at 15,000 rpm and 4◻), rinsed with 70% (v/v) EtOH and re-suspended in RNAse-free ddH_2_O. A further polysaccharide precipitation was conducted as described by (Asif, Dhawan, and Nath 2012) to remove contaminants and obtain higher quality RNA material. All RNA samples were assessed for concentration and purity using the NanoDrop ND-1000 (Thermo Fisher Scientific) and absence of RNA degradation was confirmed on a Bioanalyzer 2100 with High Sensitivity RNA chip (RNA 5000 Pico; Agilent Technologies).

### RT-qPCR

Initial validation of the flg22 treatment was assessed with real-time (RT) quantitative polymerase chain reactions (qPCR) on a QuantStudio 6 instrument (Thermo Fisher Scientific), using the total RNA extracted for the CAGE library preparations and the ΔΔC_t_ method. The list of exon/exon junction spanning primers is available in **Supplemental Dataset 10**.

### CAGE Library Construction, Filtering and Mapping

CAGE libraries were prepared as in (H. Takahashi et al. 2012) from 5 μg of total RNA and sequenced on an Illumina HiSeq 2000 platform with 30% of Phi-X spike-ins. Filtering and mapping of CAGE libraries were processed as in (Thieffry et al. 2020). Briefly, linker sequences were trimmed with FASTX Toolkit v0.0.13 (http://hannonlab.cshl.edu), and only the first 25 nt were retained. Filtering for a minimum Phred score of Q30 in 50% of the bases was applied. Clean reads were mapped on TAIR10 with Bowtie v1.1.2 (Langmead et al. 2009) and the 5’-ends of uniquely mapped reads were summed at single-bp resolution to obtain CAGE transcription start sites (CTSSs). CTSS coordinates were offset by 1 bp to account for the G-addition bias (Carninci et al. 2006).

### RNA-seq Library Construction and Analysis

Purified RNA from samples that underwent 0, 10, 30 and 60 minutes of flg22 treatment were sent to Novogene, Hong kong for preparation of polyDT-selected, unstranded, paired-end 150 bp library and sequencing on a Novaseq 6000 platform. Basecalling was conducted with Cassava (v1.8). Adapters were trimmed with Cutadapt v1.18 (Martin 2011) and clean reads were mapped on the TAIR10 reference genome with HISAT2 (Kim et al. 2019) using default parameters and keeping only concordant paired alignments. Salmon (Patro et al. 2017) was used for transcript and gene quantification and digital counts were normalized to reads per kilobase million (RPKM).

### Analysis of CAGE Tag Clusters

Most of CAGE data analyses were conducted with the CAGEfightR v1.2 package (Thodberg et al. 2019), as described in (Thieffry et al. 2020). CTSSs (genomic 5’-end positions supported by CAGE tags) with at least 1 count in a minimum of 3 libraries were retained. CAGE tag clusters (TCs) were generated by neighbour-clustering of CTSSs from the same strand with a maximum distance of 20 bp. Following quantification, CAGE TCs were further filtered for a minimum of 1 TPM in at least 3 libraries (the smallest group size in our experiment). Position of the highest signal within a TC defined the TC peak. Principal component analysis (PCA) was conducted on CAGE TCs using the scaled and centered TPM-normalized expression across libraries. CAGE TCs were annotated on the basis of the position of their peak signal against a hierarchical annotation (see **Figure 2B**, right) constructed from TAIR10 (TxDb.Athaliana.BioMart.plantsmart28, Bioconductor). Enhancer candidates were identified with the clusterBidirectionally function from CAGEfightR, using a window size of 500 bp and balance threshold of 0.95 (Bhattacharyya coefficient). Further requirements for the final set of enhancer candidates were i) a bidirectional CAGE signal existing in three or more samples, ii) overlapping with TAIR10 intergenic or intronic regions based on enhancer candidate midpoint, and iii) found in nuclear chromosomes (ChrI-V).

### GO Enrichment

Gene set enrichment analyses were conducted with the gProfileR package (Reimand et al. 2007), using all detected genes as background, and correcting resulting p-values for multiple testing with the Benjamini-Hochberg method (Benjamini et al. 2001).

### Alternative TSSs and Differential TC usage

All analyses relative to alternative TSSs used intragenic CAGE TCs contributing at least 10% to the total expression of their host gene in at least 3 libraries. Cases of differential TC usage (DTUs) in the flg22 time course were determined with the diffSplice/topSplice functions from the limma package (Ritchie et al. 2015), testing for a difference of log_2_(fold-change) of each TC compared to the average log_2_(fold-change) of all other TCs within the same gene (*t*-test). A minimum effect size of 2 in at least one time course comparison was required, and *P*-values were corrected for multiple testing with the Benjamini-Hochberg method (Benjamini et al. 2001).

### Disruption of Protein Domains

The catalogue of protein domains for *Arabidopsis thaliana* was downloaded from TAIR10 official website (www.arabidopsis.org). The position of amino acids were mapped back to the reference genome with the pmapFromTranscripts function from the GenomicFeatures Bioconductor package (Lawrence et al. 2013). Sense CAGE TCs falling within a protein domain or located downstream of a domain within the same gene were defined as domain-disruptive, using TC peak as reference position.

### Signal Peptide Analysis

Amino acid sequence of all TAIR10 proteins were retrieved from www.arabidopsis.org and scanned with SignalP-5.0 (José Juan Almagro Armenteros et al. 2019) and TargetP-2.0 (Jose Juan Almagro Armenteros et al. 2019) nd to identify localization signals in the 5’-end. Position of the predicted cleavage site was multiplied by 3 and mapped back in genomic space. First, loss of localization signal was assessed by considering CAGE TC peaks located within or downstream predicted signal peptides. In a second time, identification of relative TSS switches leading to inclusion or exclusion of signal peptide was considered on the basis of detected DTUs (see above).

### Upstream Open Reading Frame Analysis

The set of *Arabidopsis thaliana* uORFs was obtained from (Kurihara et al. 2018). After duplication and removal of inconsistent entries, uORF regions were mapped into genomic space. CAGE TCs within or downstream of uORFs were selected on the basis of their TC peak location. The only exception to this procedure were the uORFs in the *MEKK1* gene which were not contained in the Kurihara et al. (2018) dataset and which were identified by manual sequence scanning.

### Differential Expression Analyses and Clustering

Differential expression analysis was carried out with edgeR (Robinson, McCarthy, and Smyth 2010; McCarthy, Chen, and Smyth 2012) and limma (Ritchie et al. 2015) Bioconductor packages. Counts were modeled using *~ genotype * timepoint*, to capture the effects of the exosome mutants (*genotype*), the time course (*timepoint)*, and their interaction (“*” operator). The same analysis was executed at gene-level, where TC counts were aggregated when belonging to the same gene. Differentially expressed genes in the flg22 induction were selected for hierarchical clustering (with clustering method: complete, and clustering distance: correlation) on the basis of the normalized expression from the wild type samples, and visualized with the pheatmap R package.

### Transcription Factors and Enrichment of Binding Sites

AGRIS AtTFDB (Palaniswamy et al. 2006) database was used to identify genes encoding transcription factors in our experiment on the basis of AGI (Arabidopsis Genome Initiative) gene identification numbers. The position weight matrices (PWM) of each TF-encoding gene found in the flg22-responsive cluster 5 was recovered from JASPAR Plantae CORE database (Fornes et al. 2020). When no specific PWM was available, the protein sequence of the TF was used for JASPAR profile inference. A list of matrix identification numbers is available in **Supplemental Dataset 9**. Promoter regions, defined as the 500 bp stretch upstream of the closest CAGE TC peak to each annotated TAIR10 TSS, were scanned for TF motif matches with FIMO (Find Individual Motif Occurrences) (Grant, Bailey, and Noble 2011), requiring a minimum similarity of 70%. Finally, for each transcription factor in cluster 5, the genome-wide average number of matches within promoters was used as the background frequency to obtain the enrichment ratio in promoters of each DEG cluster. The resulting binding enrichments were visualized as hierarchically-clustered heatmaps.

## Data, Code, and Genome Browser Availability

CAGE libraries and processed files are available on the GEO database (ncbi.nlm.nih.gov/geo) under accessions GSE136356 and GSE143590 (access token: **khwbyaqcdzaztgj**). RNA-seq libraries and processed files are available under accession GSE144356 (access token: **khwbyaqcdzaztgj**). CAGE TCs and CAGE signal for all libraries are publicly available on the online JBrowse of www.arabidopsis.org. R scripts are publicly available on the GitHub repository athieffry/Thieffry_et_al_2021.

## Author Contributions

AT, AS and PB designed the study. JB made the CAGE libraries. AT conducted all other experiments and AB helped with total RNA extractions. AT made all analyses and figures. PB, AT and AS wrote the paper with inputs from all authors.

## List of Supplemental Datasets

Supplemental Dataset 1: CAGE TCs

Supplemental Dataset 2: Differential expression analyses

Supplemental Dataset 3: CAGE TC switches

Supplemental Dataset 4: Protein domains

Supplemental Dataset 5: Localization signals

Supplemental Dataset 6: uORFs

Supplemental Dataset 7: Enhancer candidates

Supplemental Dataset 8: DEG clusters

Supplemental Dataset 9: JASPAR matrices

Supplemental Dataset 10: Oligonucleotides

## List of Supplemental Figure

Supplemental Figure 1: PTI response marker genes in flg22 induction and differentially expressed genes

Supplemental Figure 2: CAGE-defined enhancer candidates

Supplemental Figure 3: Validation of CAGE-defined expression clusters by RNA-seq

Supplemental Figure 4: C5 transcription factors binding sites in promoter of DEG cluster genes

## Acknowledgements

Kian Hématy (*rrp4-2*), Dominique Gagliardi (*hen2-4*), are thanked for providing mutant seeds. Work in AS lab was supported by grants from Novo Nordisk Foundation and Lundbeck Foundation. Work in PB lab was supported by grants from Novo Nordisk Foundation (Hallas Møller stipend 2010), and Villum Foundation (project grant 13397).

---

**Supplemental Figure 1.**
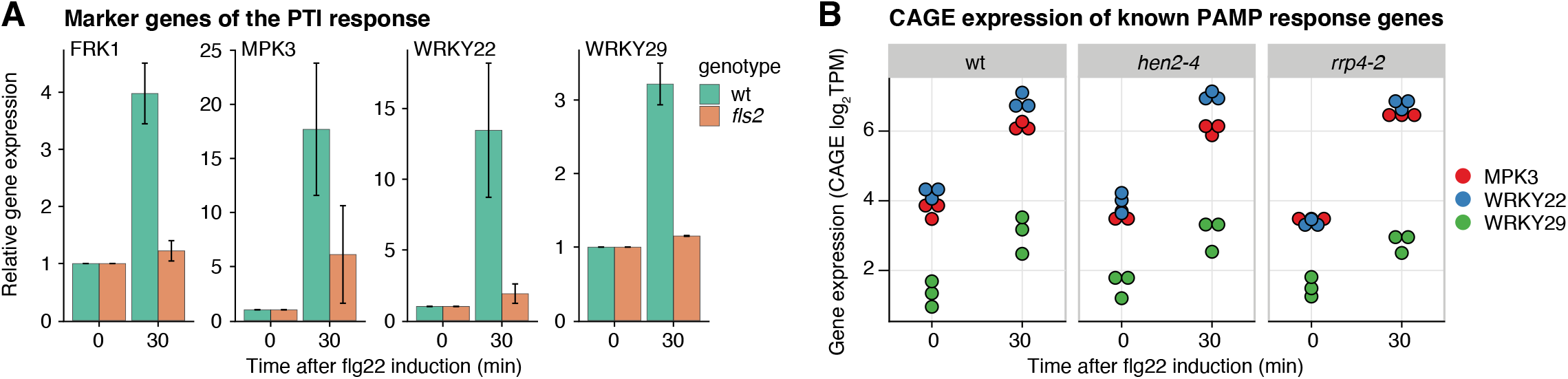
PTI response marker genes in flg22 induction and differentially expressed genes (supports Figure 1) **(A)** Relative expression of FRK1 (At2g19190), MPK3 (At3g45640), WRKY22 (At4g01250), and WRKY29 (At4g23550) as measured by real-time polymerase chain reaction at 0 and 30 minutes after flg22 treatment (X-axis). Y-axis shows expression relative to time point zero. Colors indicate genetic backgrounds and error bars indicate standard deviation. **(B)** CAGE expression of PAMP-response genes. X-axis shows time after flg22-induction, in minutes. Y-axis is the normalized CAGE expression in transcripts per millions (TPM), on log_2_ scale. Colors indicate genes and panels separate genotypes.

**Supplemental Figure 2.**
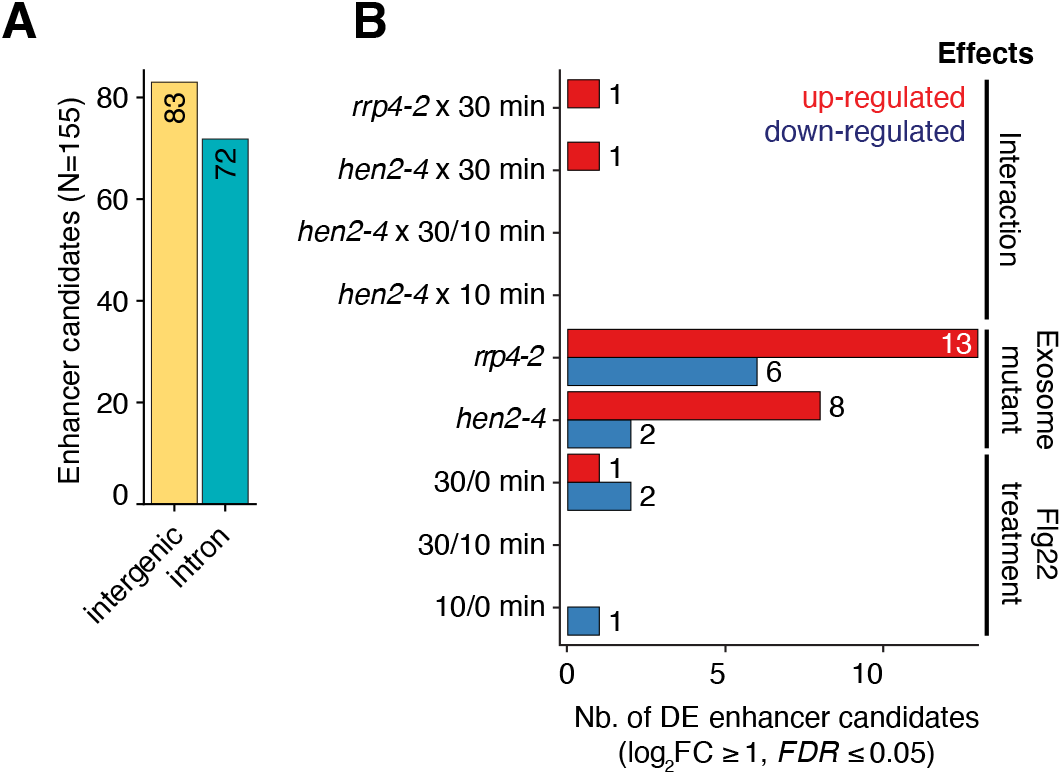
CAGE-defined enhancer candidates. **(A)** Number of enhancer candidates (Y-axis) in intergenic (yellow) or intronic regions (blue). **(B)** Differential expression analysis of enhancer candidates using CAGE expression. X-axis shows the number of enhancer candidates differentially expressed as a result of comparisons (Y-axis) which can be classified in three main effects: the flg22 treatment (indicated by minutes after treatment), exosome-related mutant genotypes (*rrp4-2*, *hen2-4*), and the interactions thereof (e.g. *rrp4-2* × 30 min). Red and blue bars indicate up- and down-regulation, respectively.

**Supplemental Figure 3.**
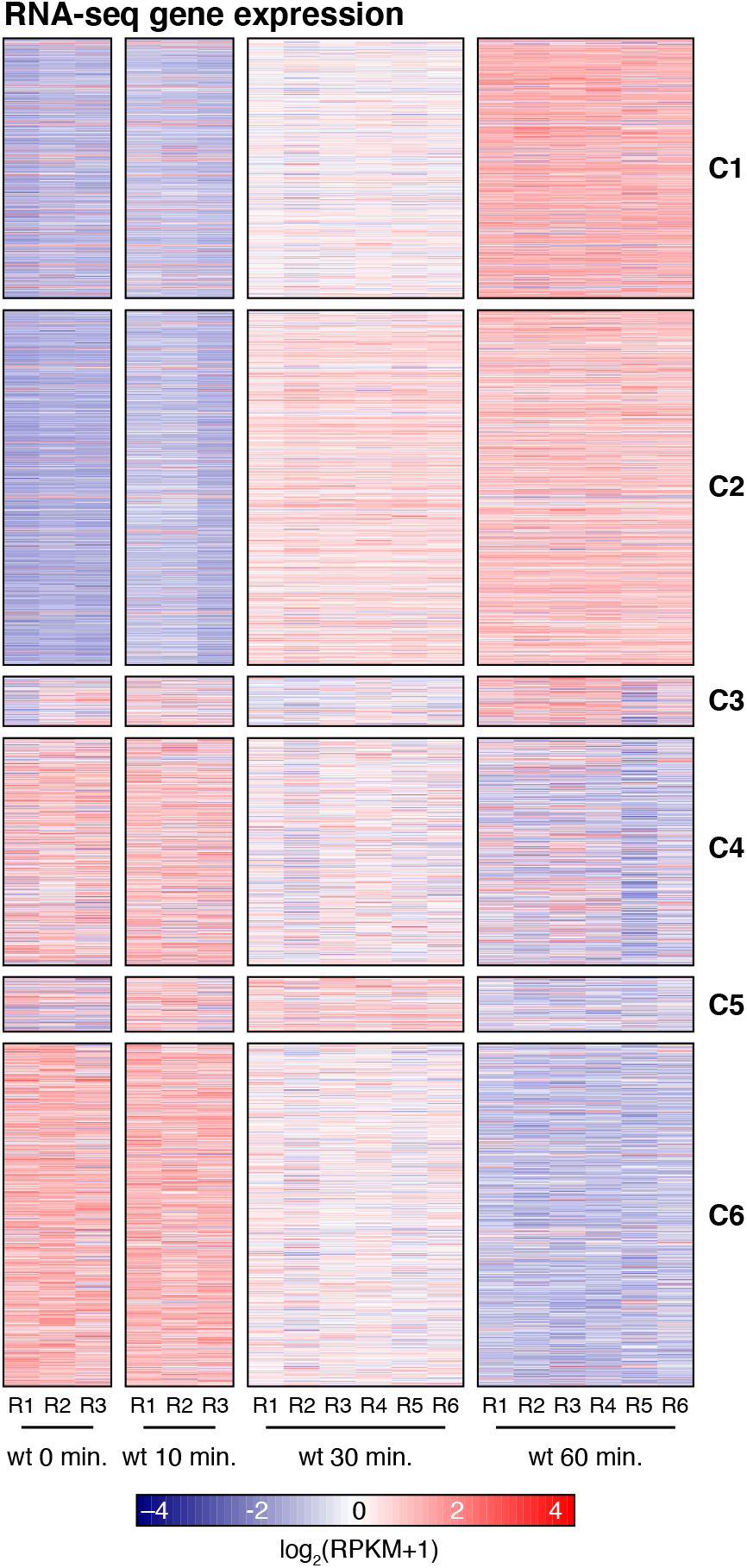
Validation of CAGE-defined expression clusters by RNA-seq (supports Figure 5) Gene expression heatmap of wild type samples treated with flg22 peptide for 0, 10, 30, and 60 minutes, as measured by RNA sequencing. Each row represents a gene, with gene order and clusters are defined by the clustering of CAGE data in Figure 5A. Columns show replicates at respective time points. Cell color indicates RPKM-normalized and log_2_-transformed RNA-seq RPKM expression with a pseudocount of 1. Labels to the right refer to CAGE-defined DEG clusters from Figure 5A.

**Supplemental Figure 4.**
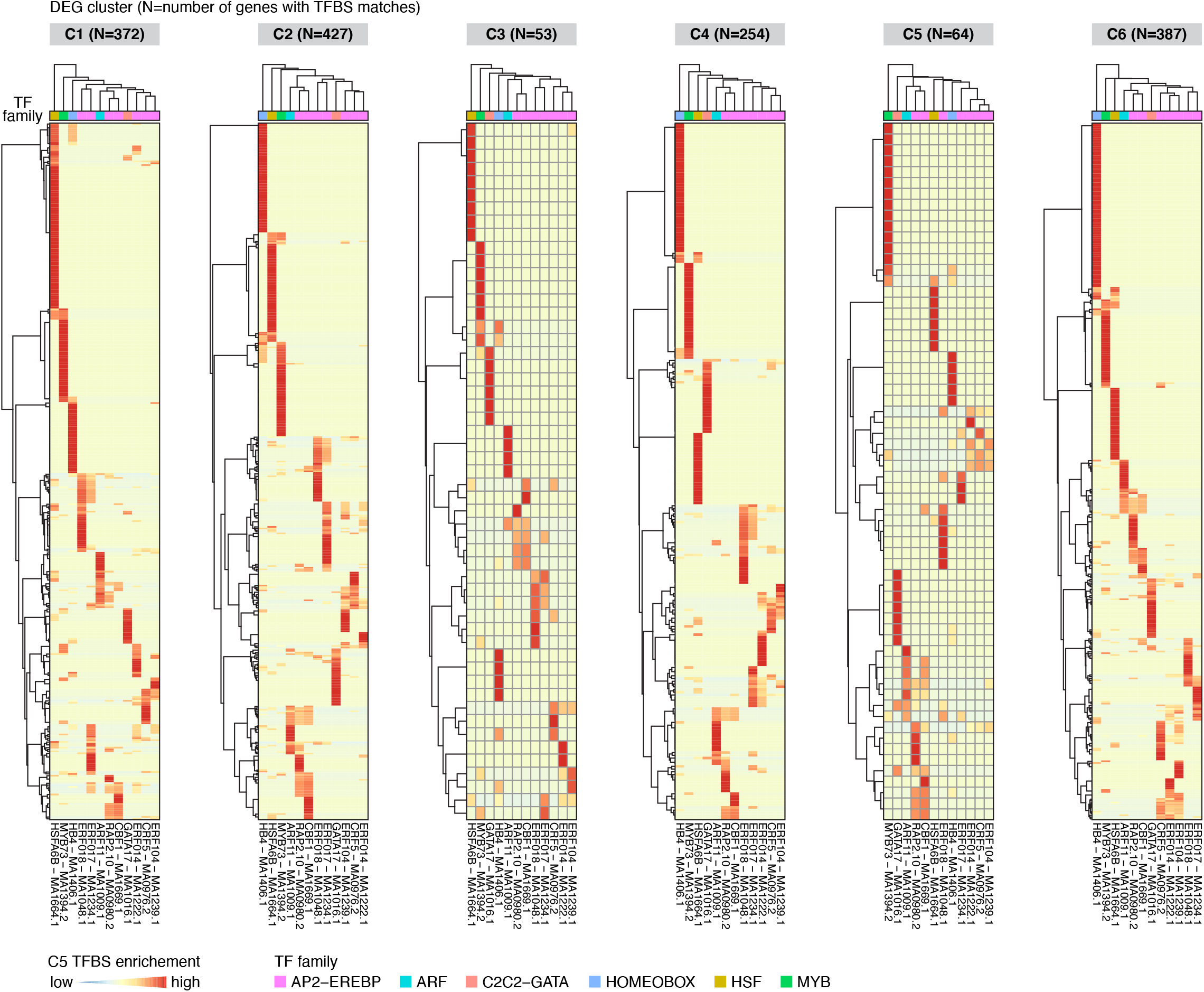
C5 transcription factors binding sites in promoter of DEG cluster genes (supports Figure 6) Enrichment of cluster 5 transcription factor binding sites in the promoter regions of genes responding to the flg22 time course. Organized as in Figure 6C, for each individual DEG cluster from Figure 5A.

## Parsed Citations

Almagro Armenteros, Jose Juan, Marco Salvatore, Olof Emanuelsson, Ole Winther, Gunnar von Heijne, Arne Elofsson, and Henrik Nielsen. 2019. “Detecting Sequence Signals in Targeting Peptides Using Deep Learning.” Life Science Alliance 2 (5). https://doi.org/10.26508/lsa.201900429. Google Scholar: Author Only Title Only Author and Title

Almagro Armenteros, José Juan, Konstantinos D. Tsirigos, Casper Kaae Sønderby, Thomas Nordahl Petersen, Ole Winther, Søren Brunak, Gunnar von Heijne, and Henrik Nielsen. 2019. “SignalP 5.0 Improves Signal Peptide Predictions Using Deep Neural Networks.” Nature Biotechnology 37 (4): 420–23. Google Scholar: Author Only Title Only Author and Title

Andersson, Robin, Claudia Gebhard, Irene Miguel-Escalada, Ilka Hoof, Jette Bornholdt, Mette Boyd, Yun Chen, et al. 2014. “An Atlas of Active Enhancers across Human Cell Types and Tissues.” Nature 507 (7493): 455–61. Google Scholar: Author Only Title Only Author and Title

Andersson, Robin, and Albin Sandelin. 2020. “Determinants of Enhancer and Promoter Activities of Regulatory Elements.” Nature Reviews. Genetics 21 (2): 71–87. Google Scholar: Author Only Title Only Author and Title

Andreasson, Erik, Thomas Jenkins, Peter Brodersen, Stephan Thorgrimsen, Nikolaj H. T. Petersen, Shijiang Zhu, Jin-Long Qiu, et al. 2005. “The MAP Kinase Substrate MKS1 Is a Regulator of Plant Defense Responses.” The EMBO Journal 24 (14): 2579–89. Google Scholar: Author Only Title Only Author and Title

Arner, Erik, Carsten O. Daub, Kristoffer Vitting-Seerup, Robin Andersson, Berit Lilje, Finn Drabløs, Andreas Lennartsson, et al. 2015. “Transcribed Enhancers Lead Waves of Coordinated Transcription in Transitioning Mammalian Cells.” Science 347 (6225): 1010–14. Google Scholar: Author Only Title Only Author and Title

Asai, Tsuneaki, Guillaume Tena, Joulia Plotnikova, Matthew R. Willmann, Wan-Ling Chiu, Lourdes Gomez-Gomez, Thomas Boller, Frederick M. Ausubel, and Jen Sheen. 2002. “MAP Kinase Signalling Cascade in Arabidopsis Innate Immunity.” Nature 415 (6875): 977–83. Google Scholar: Author Only Title Only Author and Title

Asif, Mehar H., Puneet Dhawan, and Pravendra Nath. 2012. “ASimple Procedure for the Isolation of High Quality RNAfrom Ripening Banana Fruit.” Plant Molecular Biology Reporter / ISPMB 18 (2): 109–15. Google Scholar: Author Only Title Only Author and Title

Bazin, Jeremie, Kiruthiga Mariappan, Yunhe Jiang, Thomas Blein, Ronny Voelz, Martin Crespi, and Heribert Hirt. 2020. “Role of MPK4 in Pathogen-Associated Molecular Pattern-Triggered Alternative Splicing in Arabidopsis.” PLoS Pathogens 16 (4): e1008401. Google Scholar: Author Only Title Only Author and Title

Bethke, Gerit, Tino Unthan, Joachim F. Uhrig, Yvonne Pöschl, Andrea A. Gust, Dierk Scheel, and Justin Lee. 2009. “Flg22 Regulates the Release of an Ethylene Response Factor Substrate from MAP Kinase 6 in Arabidopsis Thaliana via Ethylene Signaling.” Proceedings of the National Academy of Sciences of the United States of America 106 (19): 8067–72. Google Scholar: Author Only Title Only Author and Title

Borchardt, Erin K., Nicole M. Martinez, and Wendy V. Gilbert. 2020. “Regulation and Function of RNAPseudouridylation in Human Cells.” Annual Review of Genetics 54 (November): 309–36. Google Scholar: Author Only Title Only Author and Title

Bousquet-Antonelli, C., C. Presutti, and D. Tollervey. 2000. “Identification of a Regulated Pathwayfor Nuclear Pre-mRNATurnover.” Cell 102 (6): 765–75. Google Scholar: Author Only Title Only Author and Title

Bresson, Stefan, Alex Tuck, Desislava Staneva, and David Tollervey. 2017. “Nuclear RNADecay Pathways Aid Rapid Remodeling of Gene Expression in Yeast.” Molecular Cell 65 (5): 787–800.e5. Google Scholar: Author Only Title Only Author and Title

Carninci, Piero, Albin Sandelin, Boris Lenhard, Shintaro Katayama, Kazuro Shimokawa, Jasmina Ponjavic, Colin A. M. Semple, et al. 2006. “Genome-Wide Analysis of Mammalian Promoter Architecture and Evolution.” Nature Genetics 38 (6): 626–35. Google Scholar: Author Only Title Only Author and Title

Caro, Elena, Hume Stroud, Maxim V. C. Greenberg, Yana V. Bernatavichute, Suhua Feng, Martin Groth, Ajay A. Vashisht, James Wohlschlegel, and Steve E. Jacobsen. 2012. “The SET-Domain Protein SUVR5 Mediates H3K9me2 Deposition and Silencing at Stimulus Response Genes in a DNAMethylation–Independent Manner.” PLoS Genetics. https://doi.org/10.1371/journal.pgen.1002995. Google Scholar: Author Only Title Only Author and Title

Chen, Hsing-Yu, En-Jung Hsieh, Mei-Chun Cheng, Chien-Yu Chen, Shih-Ying Hwang, and Tsan-Piao Lin. 2016. “ORA47 (octadecanoid-Responsive AP2/ERF-Domain Transcription Factor 47) Regulates Jasmonic Acid and Abscisic Acid Biosynthesis and Signaling through Binding to a Novel Cis-Element.” The New Phytologist 211 (2): 599–613. Google Scholar: Author Only Title Only Author and Title

Chinchilla, Delphine, Zsuzsa Bauer, Martin Regenass, Thomas Boller, and Georg Felix. 2006. “The Arabidopsis Receptor Kinase FLS2 Binds flg22 and Determines the Specificityof Flagellin Perception.” The Plant Cell 18 (2): 465–76. Google Scholar: Author Only Title Only Author and Title

Chinchilla, Delphine, Cyril Zipfel, Silke Robatzek, Birgit Kemmerling, Thorsten Nürnberger, Jonathan D. G. Jones, Georg Felix, and Thomas Boller. 2007. “AFlagellin-Induced Complex of the Receptor FLS2 and BAK1 Initiates Plant Defence.” Nature 448 (7152): 497–500. Google Scholar: Author Only Title Only Author and Title

Chlebowski, Aleksander, Michał Lubas, Torben Heick Jensen, and Andrzej Dziembowski. 2013. “RNADecay Machines: The Exosome.” Biochimica et Biophysica Acta 1829 (6-7): 552–60. Google Scholar: Author Only Title Only Author and Title

Doherty, Colleen J., Heather A. Van Buskirk, Susan J. Myers, and Michael F. Thomashow. 2009. “Roles for Arabidopsis CAMTA Transcription Factors in Cold-Regulated Gene Expression and Freezing Tolerance.” The Plant Cell 21 (3): 972–84. Google Scholar: Author Only Title Only Author and Title

D’Ovidio, Renato, Benedetta Mattei, Serena Roberti, and Daniela Bellincampi. 2004. “Polygalacturonases, Polygalacturonase-Inhibiting Proteins and Pectic Oligomers in Plant-Pathogen Interactions.” Biochimica et Biophysica Acta 1696 (2): 237–44. Google Scholar: Author Only Title Only Author and Title

Etchells, J. Peter, Claire M. Provost, Laxmi Mishra, and Simon R. Turner. 2013. “WOX4 and WOX14 Act Downstreamof the PXY Receptor Kinase to Regulate Plant Vascular Proliferation Independentlyof Any Role in Vascular Organisation.” Development 140 (10): 2224–34. Google Scholar: Author Only Title Only Author and Title

Eulgem, Thomas, and Imre E. Somssich. 2007. “Networks of WRKY Transcription Factors in Defense Signaling.” Current Opinion in Plant Biology 10 (4): 366–71. Google Scholar: Author Only Title Only Author and Title

Felix, G., J. D. Duran, S. Volko, and T. Boller. 1999. “Plants Have a Sensitive Perception System for the Most Conserved Domain of Bacterial Flagellin.” The Plant Journal 18 (3): 265–76. Google Scholar: Author Only Title Only Author and Title

Fornes, Oriol, Jaime A. Castro-Mondragon, Aziz Khan, Robin van der Lee, Xi Zhang, Phillip A. Richmond, Bhavi P. Modi, et al. 2020. “JASPAR 2020: Update of the Open-Access Database of Transcription Factor Binding Profiles.” Nucleic Acids Research 48 (D1): D87–92. Google Scholar: Author Only Title Only Author and Title

Frerigmann, Henning, and Tamara Gigolashvili. 2014. “MYB34, MYB51, and MYB122 Distinctly Regulate Indolic Glucosinolate Biosynthesis in Arabidopsis Thaliana.” Molecular Plant 7 (5): 814–28. Google Scholar: Author Only Title Only Author and Title

Frerigmann, Henning, Mariola Piślewska-Bednarek, Andrea Sánchez-Vallet, Antonio Molina, Erich Glawischnig, Tamara Gigolashvili, and Paweł Bednarek. 2016. “Regulation of Pathogen-Triggered Tryptophan Metabolismin Arabidopsis Thaliana by MYB Transcription Factors and Indole Glucosinolate Conversion Products.” Molecular Plant 9 (5): 682–95. Google Scholar: Author Only Title Only Author and Title

Garcia, Mary Emily, Tim Lynch, Julian Peeters, Chris Snowden, and Ruth Finkelstein. 2008. “ASmall Plant-Specific Protein Family of ABI Five Binding Proteins (AFPs) Regulates Stress Response in Germinating Arabidopsis Seeds and Seedlings.” Plant Molecular Biology 67 (6): 643–58. Google Scholar: Author Only Title Only Author and Title

Garland, William, Itys Comet, Mengjun Wu, Aliaksandra Radzisheuskaya, Leonor Rib, Kristoffer Vitting-Seerup, Marta Lloret-Llinares, Albin Sandelin, Kristian Helin, and Torben Heick Jensen. 2019. “AFunctional Link between Nuclear RNADecayand Transcriptional Control Mediated bythe Polycomb Repressive Complex 2.” Cell Reports 29 (7): 1800–1811.e6. Google Scholar: Author Only Title Only Author and Title

Gómez-Gómez, L., and T. Boller. 2000. “FLS2: An LRR Receptor-like Kinase Involved in the Perception of the Bacterial Elicitor Flagellin in Arabidopsis.” Molecular Cell 5 (6): 1003–11. Google Scholar: Author Only Title Only Author and Title

Goralogia, Greg S., Tong-Kun Liu, Lin Zhao, Paul M. Panipinto, Evan D. Groover, Yashkarn S. Bains, and Takato Imaizumi. 2017. “CYCLING DOF FACTOR 1 Represses Transcription through the TOPLESS Co-Repressor to Control Photoperiodic Flowering in Arabidopsis.” The Plant Journal 92 (2): 244–62. Google Scholar: Author Only Title Only Author and Title

Grant, Charles E., Timothy L. Bailey, and William Stafford Noble. 2011. “FIMO: Scanning for Occurrences of a given Motif.” Bioinformatics 27 (7): 1017–18. Google Scholar: Author Only Title Only Author and Title

Guo, Meng, Jin-Hong Liu, Xiao Ma, De-Xu Luo, Zhen-Hui Gong, and Ming-Hui Lu. 2016. “The Plant Heat Stress Transcription Factors (HSFs): Structure, Regulation, and Function in Response to Abiotic Stresses.” Frontiers in Plant Science 7 (February): 114.

Hématy, Kian, Yannick Bellec, Ram Podicheti, Nathalie Bouteiller, Pauline Anne, Céline Morineau, Richard P. Haslam, et al. 2016. “The Zinc-Finger Protein SOP1 Is Required for a Subset of the Nuclear Exosome Functions in Arabidopsis.” PLoS Genetics 12 (2): e1005817. Google Scholar: Author Only Title Only Author and Title

Hickman, Richard, Marcel C. Van Verk, Anja J. H. Van Dijken, Marciel Pereira Mendes, Irene A. Vroegop-Vos, Lotte Caarls, Merel Steenbergen, et al. 2017. “Architecture and Dynamics of the Jasmonic Acid Gene Regulatory Network.” The Plant Cell 29 (9): 2086–2105. Google Scholar: Author Only Title Only Author and Title

Ichimura, Kazuya, Catarina Casais, Scott C. Peck, Kazuo Shinozaki, and Ken Shirasu. 2006. “MEKK1 Is Required for MPK4 Activation and Regulates Tissue-Specific and Temperature-Dependent Cell Death in Arabidopsis.” The Journal of Biological Chemistry 281 (48): 36969–76. Google Scholar: Author Only Title Only Author and Title

Illgen, Sylvia, Stefanie Zintl, Ellen Zuther, Dirk K. Hincha, and Thomas Schmülling. 2020. “Characterisation of the ERF102 to ERF105 Genes of Arabidopsis Thaliana and Their Role in the Response to Cold Stress.” Plant Molecular Biology 103 (3): 303–20. Google Scholar: Author Only Title Only Author and Title

Jacob, Florence, Barbara Kracher, Akira Mine, Carolin Seyfferth, Servane Blanvillain-Baufumé, Jane E. Parker, Kenichi Tsuda, Paul Schulze-Lefert, and Takaki Maekawa. 2018. “ADominant-Interfering camta3 Mutation Compromises Primary Transcriptional Outputs Mediated by Both Cell Surface and Intracellular Immune Receptors in Arabidopsis Thaliana.” The New Phytologist 217 (4): 1667–80. Google Scholar: Author Only Title Only Author and Title

Janeway, C. A., Jr. 1989. “Approaching the Asymptote? Evolution and Revolution in Immunology.” Cold Spring Harbor Symposia on Quantitative Biology 54 Pt 1: 1–13. Google Scholar: Author Only Title Only Author and Title

Kaku, Hanae, Yoko Nishizawa, Naoko Ishii-Minami, Chiharu Akimoto-Tomiyama, Naoshi Dohmae, Koji Takio, Eiichi Minami, and Naoto Shibuya. 2006. “Plant Cells Recognize Chitin Fragments for Defense Signaling through a Plasma Membrane Receptor.” Proceedings of the National Academy of Sciences of the United States of America 103 (29): 11086–91. Google Scholar: Author Only Title Only Author and Title

Kim, Daehwan, Joseph M. Paggi, Chanhee Park, Christopher Bennett, and Steven L. Salzberg. 2019. “Graph-Based Genome Alignment and Genotyping with HISAT2 and HISAT-Genotype.” Nature Biotechnology 37 (8): 907–15. Google Scholar: Author Only Title Only Author and Title

Kindgren, Peter, Maxim Ivanov, and Sebastian Marquardt. 2020. “Native Elongation Transcript Sequencing Reveals Temperature Dependent Dynamics of Nascent RNAPII Transcription in Arabidopsis.” Nucleic Acids Research 48 (5): 2332–47. Google Scholar: Author Only Title Only Author and Title

Kunze, Gernot, Cyril Zipfel, Silke Robatzek, Karsten Niehaus, Thomas Boller, and Georg Felix. 2004. “The N Terminus of Bacterial Elongation Factor Tu Elicits Innate Immunity in Arabidopsis Plants.” The Plant Cell 16 (12): 3496–3507. Google Scholar: Author Only Title Only Author and Title

Kurihara, Yukio, Yuko Makita, Mika Kawashima, Tomoya Fujita, Shintaro Iwasaki, and Minami Matsui. 2018. “Transcripts from Downstream Alternative Transcription Start Sites Evade uORF-Mediated Inhibition of Gene Expression in.” Proceedings of the National Academy of Sciences of the United States of America 115 (30): 7831–36. Google Scholar: Author Only Title Only Author and Title

Lange, Heike, François M. Sement, and Dominique Gagliardi. 2011. “MTR4, a Putative RNAHelicase and Exosome Co-Factor, Is Required for Proper rRNABiogenesis and Development in Arabidopsis Thaliana.” The Plant Journal 68 (1): 51–63. Google Scholar: Author Only Title Only Author and Title

Lange, Heike, Hélène Zuber, François M. Sement, Johana Chicher, Lauriane Kuhn, Philippe Hammann, Véronique Brunaud, et al. 2014. “The RNAHelicases AtMTR4 and HEN2 Target Specific Subsets of Nuclear Transcripts for Degradation bythe Nuclear Exosome in Arabidopsis Thaliana.” PLoS Genetics 10 (8): e1004564. Google Scholar: Author Only Title Only Author and Title

Langmead, Ben, Cole Trapnell, Mihai Pop, and Steven L. Salzberg. 2009. “Ultrafast and Memory-Efficient Alignment of Short DNA Sequences to the Human Genome.” Genome Biology 10 (3): R25.

Lawrence, Michael, Wolfgang Huber, Hervé Pagès, Patrick Aboyoun, Marc Carlson, Robert Gentleman, Martin T. Morgan, and Vincent J. Carey. 2013. “Software for Computing and Annotating Genomic Ranges.” PLoS Computational Biology 9 (8): e1003118. Google Scholar: Author Only Title Only Author and Title

Liu, Yukun, Peiyu Dang, Lixia Liu, and Chengzhong He. 2019. “Cold Acclimation bythe CBF-COR Pathwayin a Changing Climate: Lessons from Arabidopsis Thaliana.” Plant Cell Reports 38 (5): 511–19. Google Scholar: Author Only Title Only Author and Title

Lloret-Llinares, Marta, Evdoxia Karadoulama, Yun Chen, Luke A. Wojenski, Geno J. Villafano, Jette Bornholdt, Robin Andersson, Leighton Core, Albin Sandelin, and Torben Heick Jensen. 2018. “The RNAExosome Contributes to Gene Expression Regulation during Stem Cell Differentiation.” Nucleic Acids Research 46 (21): 11502–13. Google Scholar: Author Only Title Only Author and Title

Lozano, Roberto, Gregory T. Booth, Bilan Yonis Omar, Bo Li, Edward S. Buckler, John T. Lis, Jean-Luc Jannink, and Dunia Pino del Carpio. n.d. “RNAPolymerase Mapping in Plants Identifies Enhancers Enriched in Causal Variants.” https://doi.org/10.1101/376640. Google Scholar: Author Only Title Only Author and Title

Mao, Guohong, Xiangzong Meng, Yidong Liu, Zuyu Zheng, Zhixiang Chen, and Shuqun Zhang. 2011. “Phosphorylation of a WRKY Transcription Factor by Two Pathogen-Responsive MAPKs Drives Phytoalexin Biosynthesis in Arabidopsis.” The Plant Cell 23 (4): 1639–53. Google Scholar: Author Only Title Only Author and Title

Martin, Marcel. 2011. “Cutadapt Removes Adapter Sequences from High-Throughput Sequencing Reads.” EMBnet.journal. https://doi.org/10.14806/ej.17.1.200. Google Scholar: Author Only Title Only Author and Title

Mayer, Andreas, and L. Stirling Churchman. 2016. “Genome-Wide Profiling of RNAPolymerase Transcription at Nucleotide Resolution in Human Cells with Native Elongating Transcript Sequencing.” Nature Protocols 11 (4): 813–33. Google Scholar: Author Only Title Only Author and Title

McCarthy, Davis J., Yunshun Chen, and Gordon K. Smyth. 2012. “Differential Expression Analysis of Multifactor RNA-Seq Experiments with Respect to Biological Variation.” Nucleic Acids Research 40 (10): 4288–97. Google Scholar: Author Only Title Only Author and Title

Medzhitov, R., and C. A. Janeway Jr. 1997. “Innate Immunity: The Virtues of a Nonclonal System of Recognition.” Cell 91 (3): 295–98. Google Scholar: Author Only Title Only Author and Title

Mészáros, Tamás, Anne Helfer, Elizabeth Hatzimasoura, Zoltán Magyar, Liliya Serazetdinova, Gabino Rios, Viola Bardóczy, et al. 2006. “The Arabidopsis MAP Kinase Kinase MKK1 Participates in Defence Responses to the Bacterial Elicitor Flagellin.” The Plant Journal 48 (4): 485–98. Google Scholar: Author Only Title Only Author and Title

Miya, Ayako, Premkumar Albert, Tomonori Shinya, Yoshitake Desaki, Kazuya Ichimura, Ken Shirasu, Yoshihiro Narusaka, Naoto Kawakami, Hanae Kaku, and Naoto Shibuya. 2007. “CERK1, a LysM Receptor Kinase, Is Essential for Chitin Elicitor Signaling in Arabidopsis.” Proceedings of the National Academy of Sciences of the United States of America 104 (49): 19613–18. Google Scholar: Author Only Title Only Author and Title

Moore, M., Mo Vogel, and Kj Dietz. 2014. “The Acclimation Response to High Light Is Initiated within Seconds as Indicated by Upregulation of AP2/ERF Transcription Factor Network in Arabidopsis Thaliana.” Plant Signaling & Behavior 9 (10): 976479. Google Scholar: Author Only Title Only Author and Title

Moscatiello, Roberto, Paola Mariani, Dale Sanders, and Frans J. M. Maathuis. 2006. “Transcriptional Analysis of Calcium-Dependent and Calcium-Independent Signalling Pathways Induced by Oligogalacturonides.” Journal of Experimental Botany 57 (11): 2847–65. Google Scholar: Author Only Title Only Author and Title

Navarro, Lionel, Cyril Zipfel, Owen Rowland, Ingo Keller, Silke Robatzek, Thomas Boller, and Jonathan D. G. Jones. 2004. “The Transcriptional Innate Immune Response to flg22. Interplayand Overlap with Avr Gene-Dependent Defense Responses and Bacterial Pathogenesis.” Plant Physiology 135 (2): 1113–28. Google Scholar: Author Only Title Only Author and Title

Nürnberger, Thorsten, Frédéric Brunner, Birgit Kemmerling, and Lizelle Piater. 2004. “Innate Immunity in Plants and Animals: Striking Similarities and Obvious Differences.” Immunological Reviews 198 (April): 249–66. Google Scholar: Author Only Title Only Author and Title

Pajerowska-Mukhtar, Karolina M., Wei Wang, Yasuomi Tada, Nodoka Oka, Chandra L. Tucker, Jose Pedro Fonseca, and Xinnian Dong. 2012. “The HSF-like Transcription Factor TBF1 Is a Major Molecular Switch for Plant Growth-to-Defense Transition.” Current Biology: CB 22 (2): 103–12. Google Scholar: Author Only Title Only Author and Title

Palaniswamy, Saranyan K., Stephen James, Hao Sun, Rebecca S. Lamb, Ramana V. Davuluri, and Erich Grotewold. 2006. “AGRIS and AtRegNet. a Platformto Link Cis-Regulatory Elements and Transcription Factors into Regulatory Networks.” Plant Physiology 140 (3): 818–29. Google Scholar: Author Only Title Only Author and Title

Patro, Rob, Geet Duggal, Michael I. Love, Rafael A. Irizarry, and Carl Kingsford. 2017. “Salmon Provides Fast and Bias-Aware Quantification of Transcript Expression.” Nature Methods 14 (4): 417–19. Google Scholar: Author Only Title Only Author and Title

Qiu, Jin-Long, Berthe Katrine Fiil, Klaus Petersen, Henrik Bjørn Nielsen, Christopher J. Botanga, Stephan Thorgrimsen, Kristoffer Palma, et al. 2008. “Arabidopsis MAP Kinase 4 Regulates Gene Expression through Transcription Factor Release in the Nucleus.” The EMBO Journal 27 (16): 2214–21. Google Scholar: Author Only Title Only Author and Title

Ramonell, Katrina, Marta Berrocal-Lobo, Serry Koh, Jinrong Wan, Herb Edwards, Gary Stacey, and Shauna Somerville. 2005. “Loss-of-Function Mutations in Chitin Responsive Genes Show Increased Susceptibilityto the Powdery Mildew Pathogen Erysiphe Cichoracearum.” Plant Physiology 138 (2): 1027–36.

Rashotte, Aaron M., Michael G. Mason, Claire E. Hutchison, Fernando J. Ferreira, G. Eric Schaller, and Joseph J. Kieber. 2006. “A Subset of Arabidopsis AP2 Transcription Factors Mediates Cytokinin Responses in Concert with a Two-Component Pathway.” Proceedings of the National Academy of Sciences of the United States of America 103 (29): 11081–85. Google Scholar: Author Only Title Only Author and Title

Rayapuram, Naganand, Jean Bigeard, Hanna Alhoraibi, Ludovic Bonhomme, Anne-Marie Hesse, Joëlle Vinh, Heribert Hirt, and Delphine Pflieger. 2018. “Quantitative Phosphoproteomic Analysis Reveals Shared and Specific Targets of Mitogen-Activated Protein Kinases (MAPKs) MPK3, MPK4, and MPK6.” Molecular & Cellular Proteomics: MCP 17 (1): 61–80. Google Scholar: Author Only Title Only Author and Title

Reimand, Jüri, Meelis Kull, Hedi Peterson, Jaanus Hansen, and Jaak Vilo. 2007. “g:Profiler--a Web-Based Toolset for Functional Profiling of Gene Lists from Large-Scale Experiments.” Nucleic Acids Research 35 (Web Server issue): W193–200. Google Scholar: Author Only Title Only Author and Title

Ritchie, Matthew E., Belinda Phipson, Di Wu, Yifang Hu, Charity W. Law, Wei Shi, and Gordon K. Smyth. 2015. “Limma Powers Differential Expression Analyses for RNA-Sequencing and Microarray Studies.” Nucleic Acids Research 43 (7): e47.

Robinson, Mark D., Davis J. McCarthy, and Gordon K. Smyth. 2010. “edgeR: ABioconductor Package for Differential Expression Analysis of Digital Gene Expression Data.” Bioinformatics 26 (1): 139–40. Google Scholar: Author Only Title Only Author and Title

Schöffl, F., R. Prändl, and A. Reindl. 1998. “Regulation of the Heat-Shock Response.” Plant Physiology 117 (4): 1135–41. Google Scholar: Author Only Title Only Author and Title

Suarez-Rodriguez, Maria Cristina, Lori Adams-Phillips, Yidong Liu, Huachun Wang, Shih-Heng Su, Peter J. Jester, Shuqun Zhang, Andrew F. Bent, and Patrick J. Krysan. 2007. “MEKK1 Is Required for flg22-Induced MPK4 Activation in Arabidopsis Plants.” Plant Physiology 143 (2): 661–69. Google Scholar: Author Only Title Only Author and Title

Suer, Stefanie, Javier Agusti, Pablo Sanchez, Martina Schwarz, and Thomas Greb. 2011. “WOX4 Imparts Auxin Responsiveness to Cambium Cells in Arabidopsis.” The Plant Cell 23 (9): 3247–59. Google Scholar: Author Only Title Only Author and Title

Sun, Lirong, Yuxing Xu, Shenglong Bai, Xue Bai, Huijie Zhu, Huan Dong, Wei Wang, Xiaohong Zhu, Fushun Hao, and Chun-Peng Song. 2019. “Transcriptome-Wide Analysis of Pseudouridylation of mRNAand Non-Coding RNAs in Arabidopsis.” Journal of Experimental Botany 70 (19): 5089–5600. Google Scholar: Author Only Title Only Author and Title

Takagi, Momoko, Naoki Iwamoto, Yuta Kubo, Takayuki Morimoto, Hiroki Takagi, Fuminori Takahashi, Takumi Nishiuchi, et al. 2020. “Arabidopsis SMN2/HEN2, Encoding DEAD-Box RNAHelicase, Governs Proper Expression of the Resistance Gene SMN1/RPS6 and Is Involved in Dwarf, Autoimmune Phenotypes of mekk1 and mpk4 Mutants.” Plant & Cell Physiology 61 (8): 1507–16. Google Scholar: Author Only Title Only Author and Title

Takahashi, Hazuki, Timo Lassmann, Mitsuyoshi Murata, and Piero Carninci. 2012. “5′ End–centered Expression Profiling Using Cap-Analysis Gene Expression and next-Generation Sequencing.” Nature Protocols 7 (3): 542–61. Google Scholar: Author Only Title Only Author and Title

Takahashi, Naoki, Nobuo Ogita, Tomonobu Takahashi, Shoji Taniguchi, Maho Tanaka, Motoaki Seki, and Masaaki Umeda. 2019. “A Regulatory Module Controlling Stress-Induced Cell Cycle Arrest in.” eLife 8 (April). https://doi.org/10.7554/eLife.43944. Google Scholar: Author Only Title Only Author and Title

Thieffry, Axel, Maria Louisa Vigh, Jette Bornholdt, Maxim Ivanov, Peter Brodersen, and Albin Sandelin. 2020. “Characterization of Arabidopsis Thaliana Promoter Bidirectionalityand Antisense RNAs by Depletion of Nuclear RNADecay Pathways.” The Plant Cell, March. https://doi.org/10.1105/tpc.19.00815. Google Scholar: Author Only Title Only Author and Title

Thodberg, Malte, Axel Thieffry, Kristoffer Vitting-Seerup, Robin Andersson, and Albin Sandelin. 2019. “CAGEfightR: Analysis of 5’-End Data Using R/Bioconductor.” BMC Bioinformatics 20 (1): 487.

Torres, Marta de, Pedro Sanchez, Isabelle Fernandez-Delmond, and Murray Grant. 2003. “Expression Profiling of the Host Response to Bacterial Infection: The Transition from Basal to Induced Defence Responses in RPM1-Mediated Resistance.” The Plant Journal 33 (4): 665–76. Google Scholar: Author Only Title Only Author and Title

Truman, William, Marta Torres de Zabala, and Murray Grant. 2006. “Type III Effectors Orchestrate a Complex Interplaybetween Transcriptional Networks to Modify Basal Defence Responses during Pathogenesis and Resistance.” The Plant Journal 46 (1): 14–33. Google Scholar: Author Only Title Only Author and Title

Tuang, Za Khai, Zhenjiang Wu, Ye Jin, Yizhong Wang, Phyo Phyo Zin Oo, Guoxin Zuo, Huazhong Shi, and Wannian Yang. 2020. “Pst DC3000 Infection Alleviates Subsequent Freezing and Heat Injuryto Host Plants via a Salicylic Acid‐dependent Pathwayin Arabidopsis.” Plant, Cell & Environment. https://doi.org/10.1111/pce.13705.

Tuck, Alex Charles, Aneliya Rankova, Alaaddin Bulak Arpat, Luz Angelica Liechti, Daniel Hess, Vytautas Iesmantavicius, Violeta Castelo-Szekely, David Gatfield, and Marc Bühler. 2020. “Mammalian RNADecay Pathways Are Highly Specialized and Widely Linked to Translation.” Molecular Cell 77 (6): 1222–36.e13. Google Scholar: Author Only Title Only Author and Title

Tuorto, Francesca, and Frank Lyko. 2016. “Genome Recoding bytRNAModifications.” Open Biology 6 (12). https://doi.org/10.1098/rsob.160287. Google Scholar: Author Only Title Only Author and Title

Ushijima, Tomokazu, Kousuke Hanada, Eiji Gotoh, Wataru Yamori, Yutaka Kodama, Hiroyuki Tanaka, Miyako Kusano, et al. 2017. “Light Controls Protein Localization through Phytochrome-Mediated Alternative Promoter Selection.” Cell 171 (6): 1316–25.e12. Google Scholar: Author Only Title Only Author and Title

Vogel, Marc Oliver, Marten Moore, Katharina König, Pascal Pecher, Khalid Alsharafa, Justin Lee, and Karl-Josef Dietz. 2014. “Fast Retrograde Signaling in Response to High Light Involves Metabolite Export, MITOGEN-ACTIVATED PROTEIN KINASE6, and AP2/ERF Transcription Factors in Arabidopsis.” The Plant Cell 26 (3): 1151–65. Google Scholar: Author Only Title Only Author and Title

Wu, Haijun, Xiaoya Qu, Zhicheng Dong, Linjie Luo, Chen Shao, Joachim Forner, Jan U. Lohmann, et al. 2020. “WUSCHEL Triggers Innate Antiviral Immunity in Plant Stem Cells.” Science 370 (6513): 227–31. Google Scholar: Author Only Title Only Author and Title

Xiao, Wei, David Molina, Anna Wunderling, Dagmar Ripper, Joop E. M. Vermeer, and Laura Ragni. 2020. “Pluripotent Pericycle Cells Trigger Different Growth Outputs by Integrating Developmental Cues into Distinct Regulatory Modules.” Current Biology: CB 30 (22): 4384–98.e5. Google Scholar: Author Only Title Only Author and Title

Xu, Chuan, Joong-Ki Park, and Jianzhi Zhang. 2019. “Evidence That Alternative Transcriptional Initiation Is Largely Nonadaptive.” PLoS Biology 17 (3): e3000197. Google Scholar: Author Only Title Only Author and Title

Xu, Guoyong, George H. Greene, Heejin Yoo, Lijing Liu, Jorge Marqués, Jonathan Motley, and Xinnian Dong. 2017. “Global Translational Reprogramming Is a Fundamental Layer of Immune Regulation in Plants.” Nature 545 (7655): 487–90. Google Scholar: Author Only Title Only Author and Title

Xu, Rui, Yuhan Wang, Hao Zheng, Wei Lu, Changai Wu, Jinguang Huang, Kang Yan, Guodong Yang, and Chengchao Zheng. 2015. “Salt-Induced Transcription Factor MYB74 Is Regulated bythe RNA-Directed DNAMethylation Pathwayin Arabidopsis.” Journal of Experimental Botany 66 (19): 5997–6008. Google Scholar: Author Only Title Only Author and Title

Yang, Zheng-Ting, Sun-Jie Lu, Mei-Jing Wang, Dong-Ling Bi, Ling Sun, Shun-Fan Zhou, Ze-Ting Song, and Jian-Xiang Liu. 2014. “A Plasma Membrane-Tethered Transcription Factor, NAC062/ANAC062/NTL6, Mediates the Unfolded Protein Response in Arabidopsis.” The Plant Journal 79 (6): 1033–43. Google Scholar: Author Only Title Only Author and Title

Zhang, Huijuan, Yongbo Hong, Lei Huang, Dayong Li, and Fengming Song. 2016. “Arabidopsis AtERF014 Acts as a Dual Regulator That Differentially Modulates Immunity against Pseudomonas Syringae Pv. Tomato and Botrytis Cinerea.” Scientific Reports 6 (July): 30251. Google Scholar: Author Only Title Only Author and Title

Zhang, Jing, Gugan Eswaran, Juan Alonso-Serra, Melis Kucukoglu, Jiale Xiang, Weibing Yang, Annakaisa Elo, et al. 2019. “Transcriptional Regulatory Framework for Vascular Cambium Development in Arabidopsis Roots.” Nature Plants 5 (10): 1033–42. Google Scholar: Author Only Title Only Author and Title

Zhao, Zhi-Xue, Qin Feng, Peng-Qiang Liu, Xiao-Rong He, Jing-Hao Zhao, Yong-Ju Xu, Ling-Li Zhang, et al. 2021. “RPW8.1 Enhances the Ethylene-Signaling Pathwayto Feedback-Attenuate Its Mediated Cell Death and Disease Resistance in Arabidopsis.” The New Phytologist 229 (1): 516–31. Google Scholar: Author Only Title Only Author and Title

Zhu, Jiafu, Min Liu, Xiaobin Liu, and Zhicheng Dong. 2018. “RNAPolymerase II Activity Revealed by GRO-Seq and pNET-Seq in Arabidopsis.” Nature Plants 4 (12): 1112–23. Google Scholar: Author Only Title Only Author and Title

Zipfel, Cyril, Gernot Kunze, Delphine Chinchilla, Anne Caniard, Jonathan D. G. Jones, Thomas Boller, and Georg Felix. 2006. “Perception of the Bacterial PAMP EF-Tu bythe Receptor EFR Restricts Agrobacterium-Mediated Transformation.” Cell 125 (4): 749–60. Google Scholar: Author Only Title Only Author and Title

Zipfel, Cyril, Silke Robatzek, Lionel Navarro, Edward J. Oakeley, Jonathan D. G. Jones, Georg Felix, and Thomas Boller. 2004. “Bacterial Disease Resistance in Arabidopsis through Flagellin Perception.” Nature 428 (6984): 764–67. Google Scholar: Author Only Title Only Author and Title

